# Glucocorticoid-mediated Aβ and SCG10 upregulation evoke microtubule dysfunction and memory deficits

**DOI:** 10.1101/322610

**Authors:** Gee Euhn Choi, Ji Young Oh, Hyun Jik Lee, Chang Woo Chae, Jun Sung Kim, Young Hyun Jung, Ho Jae Han

## Abstract

We investigated glucocorticoid, a major risk factor of Alzheimer’s disease, promoted microtubule instability that culminates in memory deficits. Mice group exposed to corticosteroid had reduced trafficking of AMPAR1/2 and mitochondria into the synapse due to microtubule destabilization, which finally impaired cognitive function. Furthermore, cortisol reduced microtubule stability through the mitochondria glucocorticoid receptor (GR)-dependent pathway in SH-SY5Y cells. Cortisol translocated the Hsp70-bound GR into mitochondria before stimulating ER-mitochondria interaction via increasing GR-Bcl-2 complex. Subsequently, Aβ was produced since γ-secretase activity was upregulated by increased ER-mitochondria connectivity. Mitochondrial Ca^2+^ influx was also elevated due to ER-mitochondria bridging, resulting in activation of mTOR pathway. Subsequent autophagy inhibition failed to remove Aβ and led to its accumulation. Moreover, selective autophagy through ubiquitination of SCG10 was suppressed. We eventually showed that both elevated Aβ and SCG10 levels drive cells to fail trafficking AMPAR1/2 and mitochondria into the cell terminus. In conclusion, glucocorticoid regulates ER-mitochondria coupling, which evokes Aβ generation and SCG10 upregulation. Subsequent microtubule destabilization leads to memory impairment through failure of AMPAR1/2 or mitochondria transport into cell periphery.

## Introduction

Microtubule, polymeric structures composed of α- and β-tubulin heterodimer assembly, takes a pivotal role acting as the major highway for intracellular trafficking of necessary components. Notably, maintaining homeostasis in microtubule networks in neuronal cells is particularly important for strengthening synaptic connection and regulating axonal transport in neurons. Therefore, it is not surprising that microtubule dysfunction and following synaptic transport deficits are commonly observed in many neurodegenerative diseases. In Alzheimer’s disease, disease-associated factors like Aβ peptides are widely known to trigger microtubule dysfunction. For instances, reduced microtubule numbers and altered post-translational modification (PTM) of α-tubulins are observed (Brandt & Bakota, 2017). The microtubule dysfunction precedes memory impairment since neuronal cells failed to import memory-related receptors like AMPAR (α-amino-3-hydroxy-5-methyl-4-isoxazole propionic acid receptor) into synapses. Stable microtubule networks are well known to consolidate memory by promoting AMPAR endocytosis via MAP1B synthesis or the KIF5-mediated transport of AMPAR (Davidkova & Carroll, 2007, Mandal, Wei et al., 2011). Microtubule is also important for transporting mitochondria, which prefer the stable acetylated α-tubulin, into neuronal cell periphery to establish synaptic homeostasis and finally influence memory formation (Friedman, Webster et al., 2010). However, even though microtubule dysfunction represents a downstream of neurodegenerative cascades, the way in which alterations in microtubule networks induce memory impairments require further investigation. Therefore, an elucidation of the mechanism concerning pathogenesis of microtubule deficits and memory impairment is crucial for discovering the potential therapeutics of Alzheimer’s disease.

Stress, a major etiology of Alzheimer’s disease, is generally believed to induce alterations in microtubule networks through the glucocorticoid signaling pathway. The physiological level of glucocorticoid (137 – 283 nM in human serum) exerts an adaptive response to the nervous system whereas the stress-induced level of glucocorticoid (420 – 779 nM in human serum) triggers maladaptive effect such as reduced synaptic plasticity. Numerous reports have focused on the genomic effect of glucocorticoid on hyperphosphorylation of tau as a key regulator of microtubule destabilization in Alzheimer’s disease (Green, Billings et al., 2006). Recently, however, many changes in microtubule networks have been also observed such as change in the ratio of acetylated/tyrosinated α-tubulins rather than tau pathology in Alzheimer’s disease. Namely, it is important to define the various effects of glucocorticoid on microtubule dysfunction to establish a novel therapeutic approach to Alzheimer’s disease. Glucocorticoid mediates various signaling methods such as by the genomic pathway via glucocorticoid receptor (GR) binding to DNA and nongenomic pathway through glucocorticoid binding to the membrane GR. Growing evidence demonstrates that excessive glucocorticoid inhibits microtubule assembly through activating GR in rat C_6_ glioma cells (Akner, Wikstro et al., 1995) or hyperstabilizing the tubulin through nongenomic mechanism (Kershaw, Morgan et al., 2015). However, understanding of how glucocorticoid modulates microtubule dysfunction and subsequent Alzheimer’s disease remains unclear. Among the various effects, mitochondrial GR is of interest in the pathogenesis of Alzheimer’s disease since it plays a crucial role in Ca^2+^ homeostasis in mitochondria. Aberrant changes in Ca^2+^ regulation can damage the microtubule dynamics through elevated calpains (cleaving cytoskeletal proteins) and tangle formation, eventually leading to Alzheimer’s disease (Bezprozvanny & Mattson, 2008). Thus, identifying how glucocorticoid promotes microtubule dysfunction and memory impairment via regulating Ca^2+^ homeostasis is crucial for understanding molecular links between stress and Alzheimer’s disease.

In the present study, we determine the detrimental effects of glucocorticoid on microtubule networks using *in vivo* and *in vitro* models. We used male ICR mice exposed to glucocorticoid to assess the changes in microtubule dynamics and explore how glucocorticoid can affect the pathogenesis of Alzheimer’s disease. In addition, human neuroblastoma SH-SY5Y cells, widely used as neuronal disease model to investigate Alzheimer’s disease, were utilized to investigate the detailed mechanism of microtubule dysfunction via ER-mitochondria interaction by glucocorticoid. Overall, we demonstrate the mechanism of glucocorticoid on microtubule dysfunction and memory deficits in both suggested *in vitro* and *in vivo* models.

## Results

### The effect of corticosterone on memory impairment *in vivo*

We first examined microtubule dynamics in hippocampus of male ICR mice treated with corticosterone, the major form of glucocorticoid in rodents. Microtubule dynamics can be controlled by the intrinsic GTPase activity of tubulins and various PTMs that occur on the C-terminal tails, interacting with motor proteins and microtubule-associated proteins (Uchida & Shumyatsky, 2015). Acetylated or detysoinated α-tubulin is the marker of stable tubulin reducing microtubule depolymerization whereas tyrosinated α-tubulin is the labile tubulin. The ratio of acetylated to tyrosinated α-tubulin (A/T ratio) has been used to evaluate microtubule stability (Uchida, Martel et al., 2014). Both immunoblotting and immunofluorescent results revealed that corticosterone reduced the A/T ratio in hippocampus (Fig. 1A - 1B). Microtubule destabilization usually exerts decreased transport of necessary components into cell periphery. Thus, a failure to traffick AMPAR1/2 into synapse was shown in hippocampus of mice with corticosterone (Fig. 1C). In addition, the perinuclear clumping of mitochondria was observed in mice with corticosterone (Fig. 1D), which is the representative phenomenon for microtubule dysfunction. Since mitochondria trafficking to cell terminus was impaired, cell death would likely follow. We identified the cell viability using the TUNEL assay where the fluorescence of Alexa Fluor^TM^ 488 dye detects fragmented DNA visualization. The results showed increased cell apoptosis with corticosterone (Fig. 1E). With decreased AMPAR and mitochondria transport, the spontaneous alteration percentage for evaluating cognitive function was decreased in mice with corticosterone (Fig. 1F).

**Figure 1.**
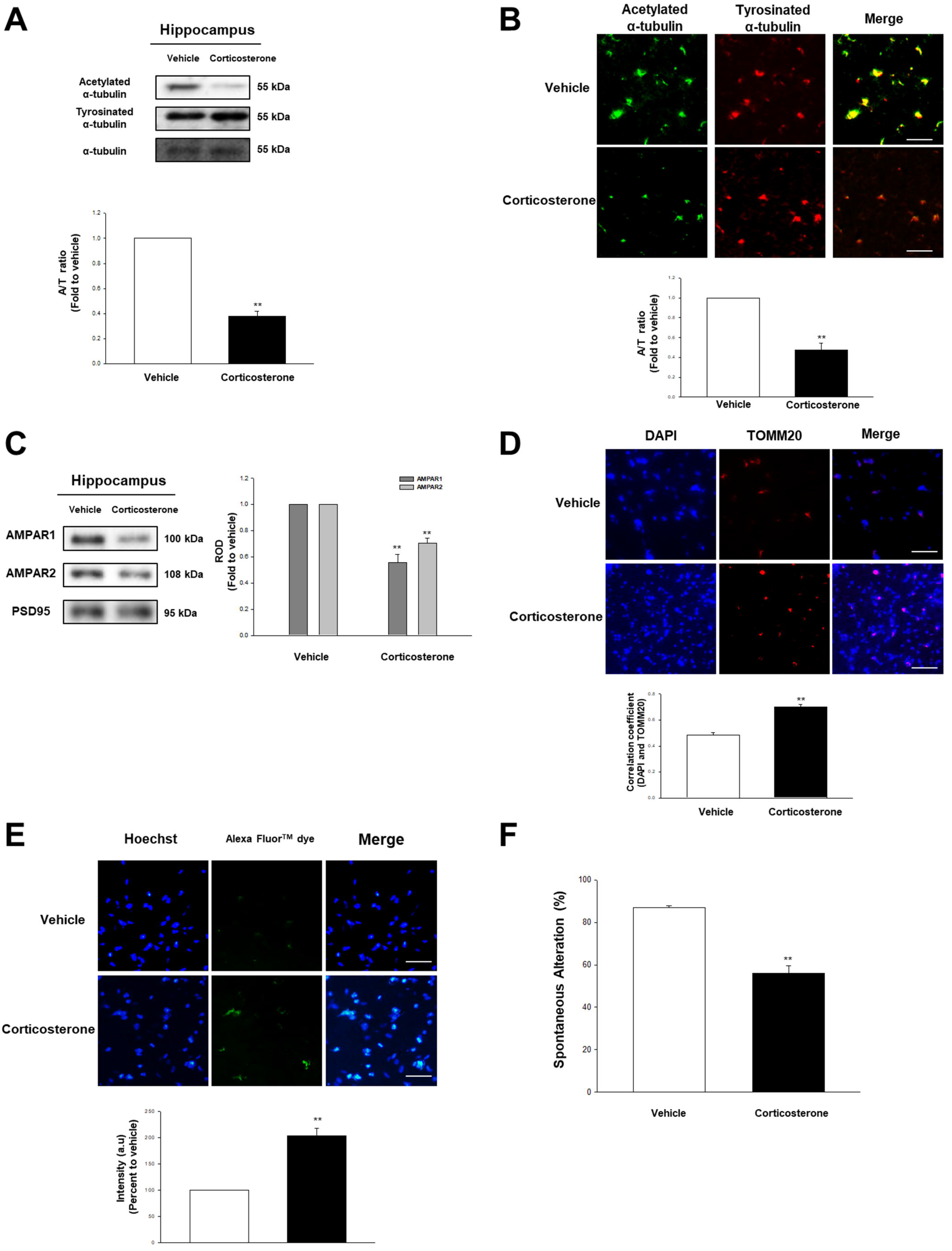
The effect of corticosterone on microtubule destabilization in the male ICR mice. (**A**) The hippocampus of male ICR mice exposed to vehicle or corticosterone (10 mg/kg) was collected. Acetylated α-tubulin, tyrosinated α-tubulin, and α-tubulin were detected by western blot. Data are reported as a mean ± SE of five independent experiments. indicates *p<0.01* versus vehicle. (**B**) Slide samples for immunohistochemistry (IHC) were immunostained with acetylated α-tubulin (green) and tyrosinated α-tubulin (red). Scale bars, 200 μm (magnification, × 200). The A/T ratio was measured and the data are reported as a mean ± SE of five independent experiments. ** indicates *p<0.01* versus vehicle. (**C**) The synaptic protein was extracted from the hippocampus of mice treated with vehicle or corticosterone (10 mg/kg). Synaptic protein expressions were normalized by loading control of synaptosome, PSD95, in western blotting results. Data are reported as a mean ± SE of five independent experiments. ** indicates *p<0.01* versus vehicle. (**D**) Slide samples for IHC were immunostained with DAPI (blue) and TOMM20 (red). Scale bars, 200 μm (magnification, × 200). Correlation coefficient analysis between DAPI and TOMM20 was done. Data are reported as a mean ± SE of five independent experiments. ** indicates *p<0.01* versus vehicle. (**E**) TUNEL assay was performed using slide samples of hippocampus from mice with vehicle or corticosterone (10 mg/kg). The intensity of green fluorescence indicates the amount of neuronal cell death. Scale bars, 200 μm (magnification, × 200). Data are reported as a mean ± SE of five independent experiments. ** indicates *p<0.01* versus vehicle. (**F**) The mice exposed to vehicle or corticosterone (10 mg/kg) were subjected to Y-maze test to evaluate memory function. Data are reported as a mean ± SE of six independent experiments. indicates *p<0.01* versus vehicle.

### Translocation of GR into mitochondria induced ER-mitochondria tethering

We evaluated the effect of cortisol, the major form of glucocorticoid in humans, on microtubule destabilization in SH-SY5Y cell lines based on *in vivo* experiments. We showed that cortisol decreased the A/T ratio of cells in a concentration and time dependent manner (Fig. 2A - 2B). Reduced A/T ratio was also detected in the immunostaining results upon cortisol (Fig. 2C). Cortisol can trigger various signaling methods like genomic pathway, nongenomic pathway, and mitochondria GR-mediated pathway. To differentiate which signaling induces the change in microtubule dynamics, actinomycin D (the nuclear transcription inhibitor) that blocks the genomic pathway of cortisol, and the cortisol-BSA, which only induces the membrane GR-mediated pathway, were treated. We found that there was no recovered A/T ratio with actinomycin D pretreatment and no significant changes in the A/T ratio with cortisol-BSA treatment (Fig EV. 1A - 1B). Co-localization of GR and COX IV, the mitochondrial marker, increased upon cortisol for 2 h (Fig. 2D). Subcellular fraction results also indicated that cortisol induced GR translocation into mitochondria, inhibited by RU 486 (competitive inhibitor of GR, Fig EV. 1C). The inactive form of GR binds to many chaperone proteins. The active form of GR detaches from heat shock protein90 (Hsp90) with ligand and is subsequently translocated into nucleus or mitochondria. Some GRs do not detach from Hsp70 or selectively associate with un-bound cytosolic Hsp70; all of which guide the translocation of GR into mitochondria (Du, McEwen et al., 2009a). Consistent with previous research, cortisol promoted GR-Hsp70 coupling while GR-Hsp90 interaction was reduced (Fig. 2E). The immunoprecipitation of mitochondrial parts also revealed that GR-Hsp70 binding was elevated when exposed to cortisol (Fig. 2F). The increased GR translocation into mitochondria was decreased with VER 155008 (Hsp70 inhibitor) pretreatment (Fig. 2G). We also found that the reduced A/T ratio was recovered with VER 155008 pretreatment (Fig EV. 1D), suggesting that mitochondrial GR regulates microtubule dynamics.

**Figure 2.**
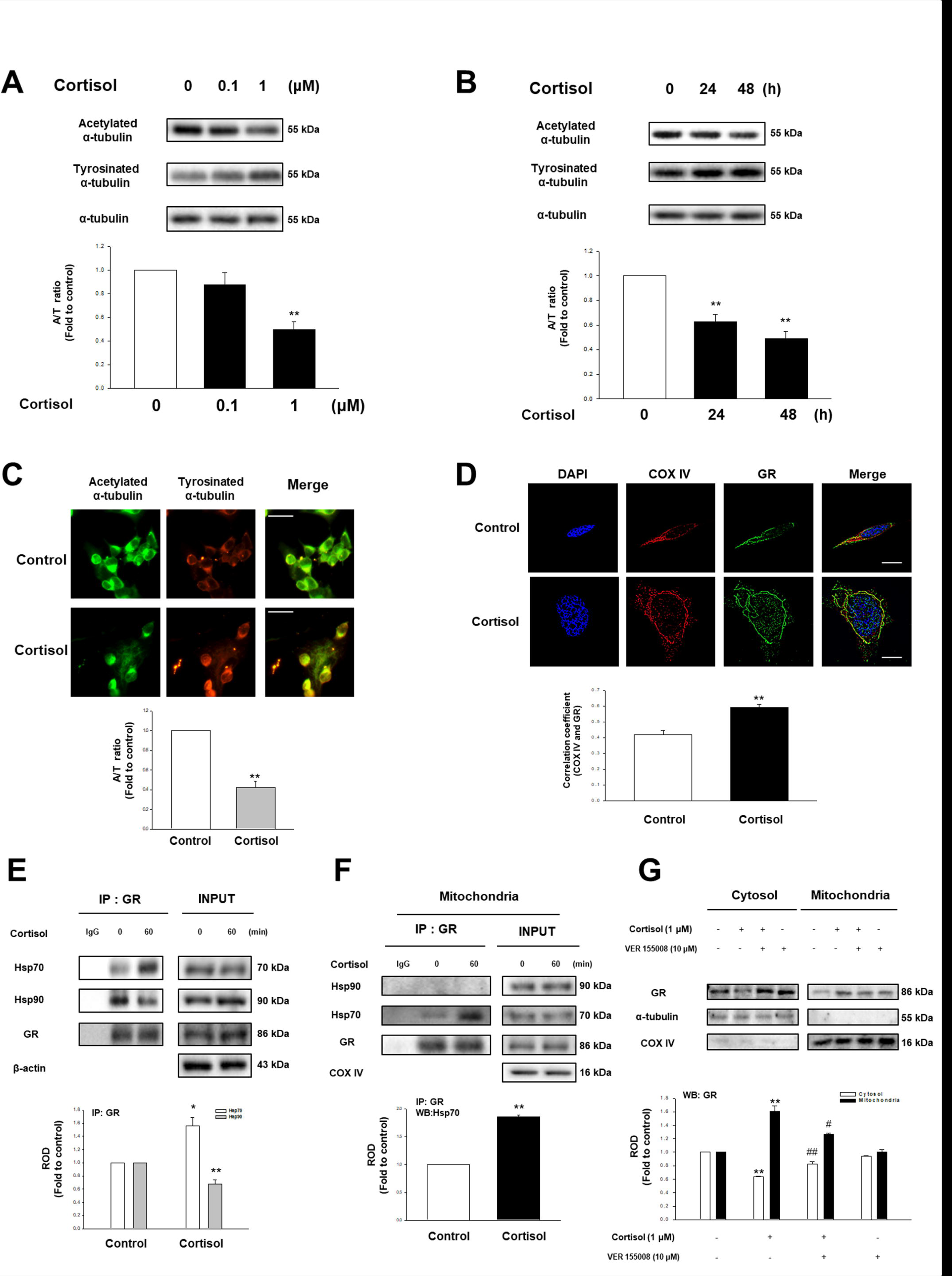
The effect of cortisol on microtubule dysfunction via GR trafficking into mitochondria in SH-SY5Y. (**A**) Cortisol (0 - 1 μM) was treated for 48 h in SH-SY5Y cells. Then, acetylated α-tubulin, tyrosinated α-tubulin, and α-tubulin were detected by western blot. Data are reported as a mean ± SE of five independent experiments. ** indicates *p<0.01* versus control. (**B**) Total cell lysates in a time response with 1 μM cortisol were subjected to western blot. Acetylated α-tubulin, tyrosinated α-tubulin, and α-tubulin were detected. Data are reported as a mean ± SE of five independent experiments. ** indicates *p<0.01* versus control. (**C**) Immunostaining of cells treated with cortisol for 48 h were visualized by Eclipse Ts2™ fluorescence microscopy. The green indicates acetylated α-tubulin and the red indicates tyrosinated α-tubulin. Data are reported as a mean ± SE of ive independent experiments. Scale bars, 100 μm (magnification, × 400). (**D**) The cells were treated with cortisol (1 μM) for 2 h which were immnunostained with DAPI (blue), COX IV (red) and glucocorticoid receptor (GR, green). The images were acquired by super-resolution radial fluctuations (SRRF) imaging system. Data are reported as a mean ± SE of five independent experiments. ** indicates *p<0.01* versus control. Scale bars, 20 μm (magnification, × 1,000). (**E**) The cells were incubated with cortisol (1 μM) for 60 min and then harvested. GR was co-immunoprecipitated with anti-Hsp70 and -Hsp90 antibodies (the left side). Expression of Hsp70, Hsp90, GR, and β-actin in total cell lysates is shown in the right side. Data are reported as a mean ± SE of four independent experiments. *,** indicates *p<0.05, p<0.01* versus control, respectively. (**F**) The mitochondrial parts of the cells treated with cortisol (1 μM) for 60 min underwent immunoprecipiatation with GR. Data are reported as a mean ± SE of five independent experiments. ** indicates *p<0.01* versus control. (**G**) The cells were incubated with VER 155008 (10 μM) for 30 min before cortisol treatment (1 μM) for 2 h. GR, α-tubulin, and COX IV were detected. Cytosolic and mitochondrial protein expressions were normalized by α-tubulin and COX IV, respectively, in western blotting results. Data are reported as a mean ± SE of four independent experiments. ** indicates *p<0.01* versus control. ^#^, ^##^ indicates *p<0.05, p<0.01* versus cortisol, respectively.

GR regulates mitochondrial calcium by binding to Bcl-2, but underlying mechanism is not understood. One of major factors in facilitating the uptake of Ca^2+^ by mitochondria is the contact between ER and mitochondria where inositol 1,4,5-triphosphate receptor (IP3R) and voltage-dependent anion-selective channel 1 (VDAC1) bridge. Thus, we examined the effect of cortisol on ER-mitochondria coupling via GR binding to Bcl-2. We found the interaction between mitochondrial protein, TOMM20, and IP3R [one of MAM (mitochondria-associated membrane of ER) proteins] was increased with cortisol treatment while the Bcl-2 was also translocated into mitochondria (Fig. 3A). We also showed that the binding between GR to Bcl-2, IP3R, and TOMM20 was increased with cortisol (Fig. 3B), suggesting that GR-Bcl-2 complex by cortisol can increase ER-mitochondria contact. As predicted, co-localization between IP3R and VDAC1 was strongly stimulated with cortisol treatment, which was reduced by RU 486 (Fig. 3C). PLA results also showed that cortisol led to IP3R-VDAC1 interactions, which were decreased with the knockdown of *bcl-2* (Fig. 3D). We then assessed protein from which each organelle ligates to form ER-mitochondria coupling. Cortisol triggered the binding between mfn2 and phosphofurin acidic cluster sorting protein 2 (PACS2), which was reversed by knockdown of *bcl-2* (Fig. 3E). Meanwhile, this had no effect on vesicle-associated protein B (VAPB)-protein tyrosine phosphatase interacting protein 51 (PTPIP51) complex (Fig EV. 2A). Similar to *in vitro* results, the hippocampus of mice with corticosterone showed increased IP3R-VDAC1 interaction binding to GR and Bcl-2 (Fig. 3F). Furthermore, we also found that ER-mitochondria contact was upregulated in hippocampus of mice exposed to corticosterone in PLA results (Fig. 3G) via mfn2-PACS2 binding (Fig. 3H – 3I). Inhibiting ER-mitochondria tethering with knockdown of *bcl-2* or RU 486 also evoked microtubule destabilization indicating that this connectivity regulates microtubule dynamics (Fig EV. 2B - 2C).

**Figure 3.**
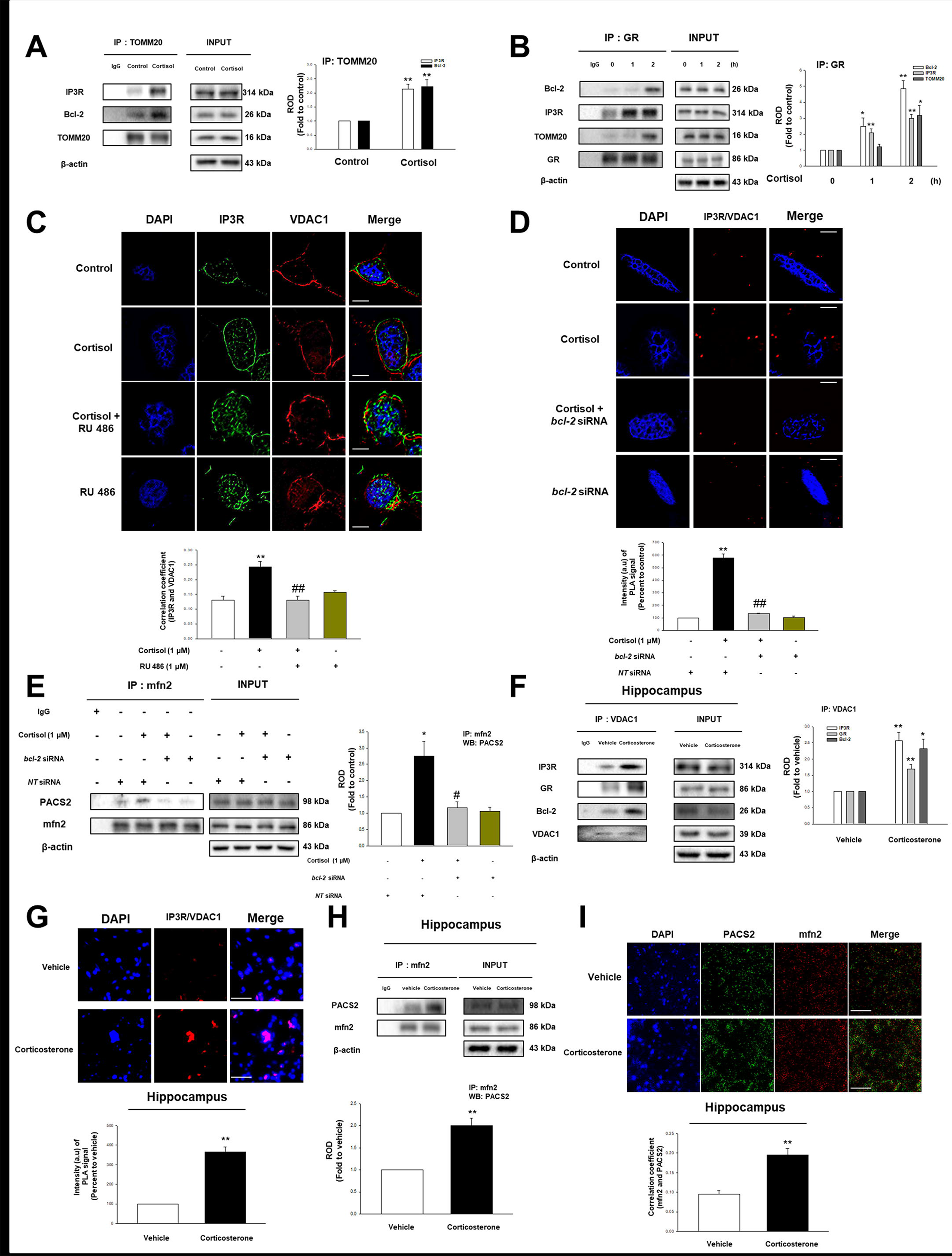
The effect of cortisol on ER-mitochondria contact via interaction between GR and Bcl-2. (**A**) The cells were incubated with cortisol (1 μM) for 2 h and then harvested. TOMM20 was co-immunoprecipitated with anti-IP3R and -Bcl-2 antibodies (the left side). Expression of IP3R, Bcl-2, TOMM20, and β-actin in total cell lysates is shown in the right side. Data are reported as a mean ± SE of four independent experiments. ** indicates *p<0.01* versus control. (**B**) The cells were incubated with cortisol (1 μM) for various time and then harvested. GR was co-immunoprecipitated with anti-Bcl-2, -IP3R and -TOMM20 antibodies (the left side). Expression of Bcl-2, IP3R, TOMM20, GR, and β-actin in total cell lysates is shown in the right side. Data are reported as a mean ± SE of four independent experiments. *,** indicates *p<0.05, p<0.01* versus control, respectively. (**C**) The cells were incubated with RU 486 (1 μM) for 30 min before cortisol treatment (1 μM) for 2 h. Co-localization of IP3R (green) and VDAC1 (red) was visualized with SRRF imaging system. DAPI was used for nuclear counterstaining (blue). Data are reported as a mean ± SE of five independent experiments. ** indicates *p<0.01* versus control and ^##^ indicates *p<0.01* versus cortisol alone. Scale bars represent 20 μm (magnification, × 900). (**D**) Knockdown of *bcl-2* was done using siRNA transfection for 24 h and then cells were treated with cortisol (1 μM) during 2 h. The cells underwent proximal ligation assay (PLA) and the red fluorescence indicates the co-localization between IP3R and VDAC1. DAPI was used for nuclear counterstaining (blue). Data are acquired by SRRF imaging system and reported as a mean ± SE of four independent experiments. ** indicates *p<0.01* versus control and ^##^ indicates *p<0.01* versus cortisol alone. Scale bars represent 20 μm (magnification, × 900). (**E**) Knockdown of *bcl-2* was done using siRNA transfection for 24 h and then cells were treated with cortisol (1 μM) during 2 h. mfn2 was co-immunoprecipitated with an anti-PACS2 antibody (the left side). Expression of PACS2, mfn2, and β-actin in total cell lysates is shown in the right side. Data are reported as a mean ± SE of four independent experiments. * indicates *p<0.05* versus control and ^#^ indicates *p<0.05* versus cortisol alone. (**F**) The hippocampus of mice exposed to vehicle or corticosterone (10 mg/kg) for 2 h was collected and lysed. VDAC1 was co-immunoprecipitated with anti-IP3R, -GR, and -Bcl-2 antibodies (the left side). Expression of IP3R, GR, Bcl-2, VDAC1, and β-actin in total cell lysates is shown in the right side. Data are reported as a mean ± SE of five independent experiments. *, ** indicates *p<0.05, p<0.01* versus vehicle, respectively. (**G**) Slide samples for IHC of mice with vehicle or corticosterone (10 mg/kg) for 2 h underwent PLA and the red fluorescence indicates the co-localization between IP3R and VDAC1. DAPI was used for nuclear counterstaining (blue). Data are reported as a mean ± SE of five independent experiments. ** indicates *p<0.01* versus vehicle. Scale bars, 200 μm (magnification, × 200). (**H**) The hippocampus of mice exposed to vehicle or corticosterone (10 mg/kg) for 2 h was collected and lysed. mfn2 was co-immunoprecipitated with an anti-PACS2 antibody (the left side). Expression of PACS2, mfn2, and β-actin in total cell lysates is shown in the right side. Data are reported as a mean ± SE of five independent experiments. ** indicates *p<0.01* versus vehicle. (**I**) Slide samples for IHC of mice with vehicle or corticosterone (10 mg/kg) for 2 h were immunostained with PACS2 (green) and mfn2 (red). DAPI was used for nuclear counterstaining (blue). Scale bars, 200 μm (magnification, × 200). Data are reported as a mean ± SE of five independent experiments. ** indicates *p<0.01* versus vehicle.

### Involvement of ER-mitochondria connectivity in Aβ formation and SCG10 upregulation

With respect to the fact that ER-mitochondria tethering evokes many physiological changes, we demonstrated that this connectivity promotes Aβ accumulation and superior cervical ganglion-10 (SCG10) upregulation. Increased ER-mitochondria connectivity is likely to upregulate Aβ level since γ-secretase localizes in ER-mitochondria contact site. We demonstrated that cortisol increased the localization of PSEN1 and C99 in the MAM marker, VAPB, which is reduced by knockdown of *bcl-2*, pretreatment of RU 486, and DAPT (γ-secretase inhibitor) via immunoprecipitation (Fig. 4A - 4B). Immunofluorescent results also suggested that both PSEN1 and C99 interact with the ER-mitochondria contact site, inhibited by knockdown of *bcl-2* or pretreatment of RU 486 and DAPT (Fig. 4C - 4F). Furthermore, the upregulated Aβ level was concentrated on ER-mitochondria contact site, also decreased by knockdown of *bcl-2*, pretreatment of RU 486, and DAPT in the immunofluorescence results (Fig. 4G - 4H). Aβ produced at MAM-mitochondria contact sites is known to be translocated into mitochondria and further moves to the extracellular compartment. Therefore, we showed that cortisol triggers the accumulation of Aβ into mitochondria, the effect of which was abolished with knockdown of *bcl-2*, pretreatment of RU 486, and DAPT, γ-secretase inhibitor (Fig. 4I). Subsequently, Aβ level was dramatically increased in the medium while it was decreased by inhibiting ER-mitochondria coupling via *bcl-2* knockdown and pretreatment of RU 486, or γ-secretase activity (Fig. 4J - 4K). Therefore, the data suggests that an increase in ER-mitochondria contact via GR and Bcl-2 coupling eventually induced amyloidosis at MAM-mitochondria contact sites.

**Figure 4.**
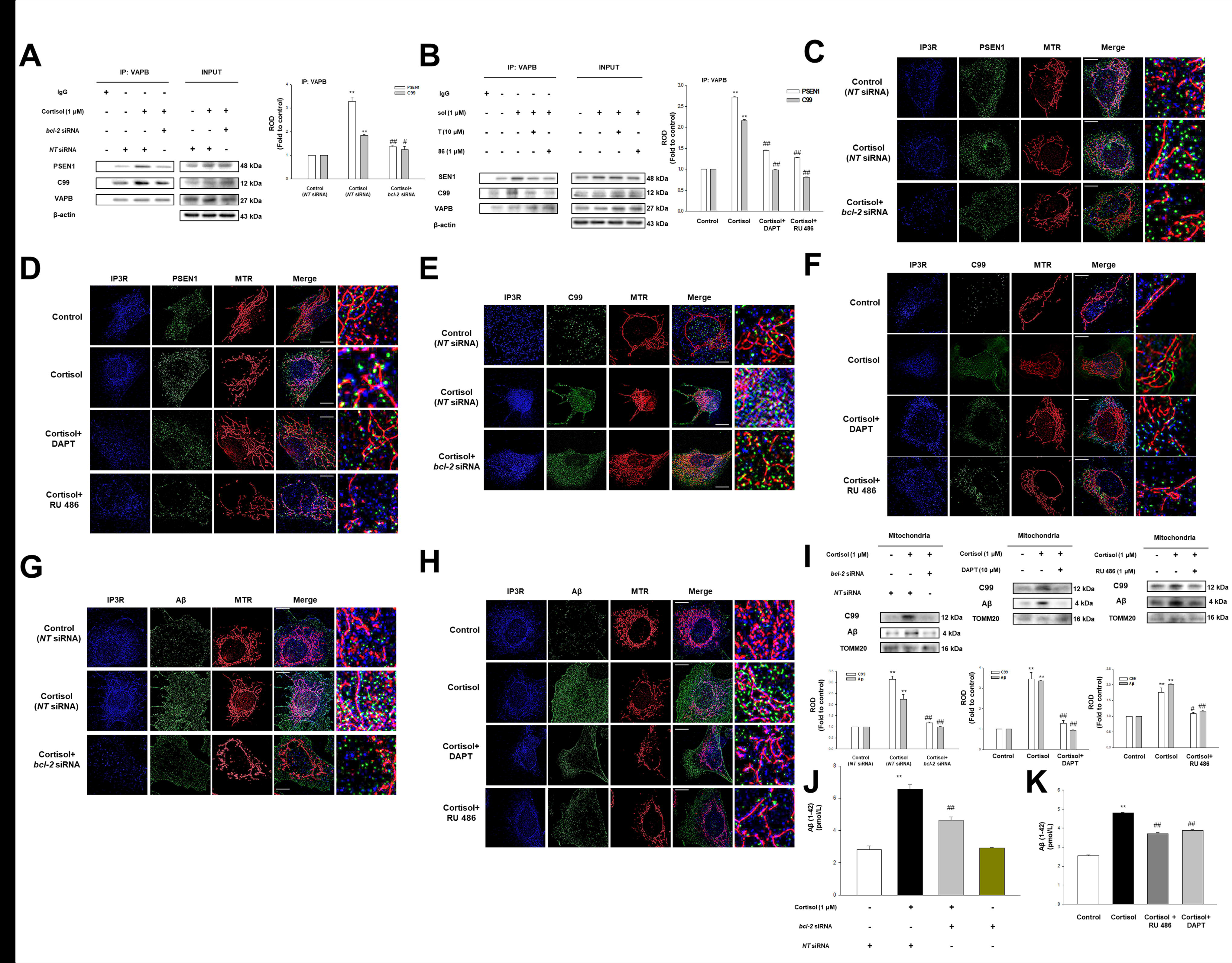
Cortisol-mediated ER-mitochondria tethering induced Aβ formation. (**A**) Knockdown of *bcl-2* was done using siRNA transfection for 24 h and then cells were treated with cortisol (1 μM) during 12 h. VAPB was co-immunoprecipitated with anti-presenilin1 (PSEN1) and -C99 antibodies (the left side). Expression of PSEN1, C99, VAPB, and β-actin in total cell lysates is shown in the right side. Data are reported as a mean ± SE of four independent experiments. ** indicates *p<0.01* versus control, respectively. ^#, ##^ indicates *p<0.05, p<0.01* versus cortisol alone, respectively. (**B**) The cells were pre-incubated with RU 486 (1 μM) or DAPT (10 μM) for 30 min before cortisol (1 μM) for 12 h. VAPB was co-immunoprecipitated with anti-PSEN1 and -C99 antibodies (the left side). Expression of PSEN1, C99, VAPB, and β-actin in total cell lysates is shown in the right side. Data are reported as a mean ± SE of four independent experiments. ** indicates *p<0.01* versus control and ^##^ indicates *p<0.01* versus cortisol alone. (**C**) Knockdown of *bcl-2* was done using siRNA transfection for 24 h and then cells were treated with cortisol (1 μM) during 12 h. Co-localization of IP3R (blue), PSEN1 (green), and Mitotracker Red (MTR, red) was visualized with SRRF imaging system. Scale bars represent 20 μm (magnification, × 1,000). n=4. (**D**) The cells were pre-incubated with RU 486 (1 μM) or DAPT (10 μM) for 30 min before cortisol (1 μM) for 12 h. Co-localization of IP3R (blue), PSEN1 (green), and MTR (red) was visualized with SRRF imaging system. Scale bars represent 20 μm (magnification, × 1,000). n=4. (**E**) Knockdown of *bcl-2* was done using siRNA transfection for 24 h and then cells were treated with cortisol (1 μM) during 12 h. Co-localization of IP3R (blue), C99 (green), and MTR (red) was visualized with SRRF imaging system. Scale bars represent 20 μm (magnification, × 1,000). n=4. (**F**) The cells were pre-incubated with RU 486 (1 μM) or DAPT (10 μM) for 30 min before cortisol (1 μM) for 12 h. Co-localization of IP3R (blue), C99 (green), and MTR (red) was visualized with SRRF imaging system. Scale bars represent 20 μm (magnification, × 1,000). n=4. (**G**) Knockdown of *bcl-2* was done using siRNA transfection for 24 h and then cells were treated with cortisol (1 μM) during 24 h. Co-localization of IP3R (blue), Aβ (green), and MTR (red) was visualized with SRRF imaging system. Scale bars represent 20 μm (magnification, × 1,000). n=4. (**H**) The cells were pre-incubated with RU 486 (1 μM) or DAPT (10 μM) for 30 min before cortisol (1 μM) for 24 h. Co-localization of IP3R (blue), Aβ (green), and MTR (red) was visualized with SRRF imaging system. Scale bars represent 20 μm (magnification, × 1,000). n=4. (**I**) Mitochondrial parts were isolated from the cells transfected with *bcl-2* siRNA (left panel) or pretreated with DAPT (10 μM, middle panel), RU 486 (1 μM, right panel) before cortisol (1 μM) during 24 h. C99, Aβ, and TOMM20 were detected. Mitochondrial protein expressions were normalized by TOMM20 in western blotting results. Data are reported as a mean ± SE of four independent experiments. ** indicates *p<0.01* versus control. ^#^, ^##^ indicates *p<0.05, p<0.01* versus cortisol, respectively. (**J**) Knockdown of *bcl-2* was done using siRNA transfection for 24 h and then cells were treated with cortisol (1 μM) during 48 h. The secreted Aβ in conditioned medium was detected using ELISA kit. Data are reported as a mean ± SE of four independent experiments. ** indicates *p<0.01* versus control and ^##^ indicates *p<0.01* versus cortisol alone. (**K**) The cells were pre-incubated with RU 486 (1 μM) or DAPT (10 μM) for 30 min before cortisol (1 μM) for 48 h. The secreted Aβ in conditioned medium was detected using ELISA kit. Data are reported as a mean ± SE of four independent experiments. ** indicates *p<0.01* versus control and ^##^ indicates *p<0.01* versus cortisol alone.

ER-mitochondria contact by cortisol also increased immunofluorescence intensity of rhod-2 which binds to mitochondrial Ca^2+^ (Fig. 5A). This indicates that Ca^2+^ was transferred from the IP3R-VDAC1 bridge, which mainly functions as a Ca^2+^ pathway from ER to mitochondria (Hedskog, Pinho et al., 2013). Induction of mitochondrial Ca^2+^ was downregulated by the knockdown of *bcl-2*, xestospongin C (the potent membrane permeable IP3R antagonist), and ruthenium red - VDAC1 inhibitor (Fig EV. 3A – 3C). Consistent Ca^2+^ transfer from ER to mitochondria is basically required to maintain ample NADH production (Cárdenas, Miller et al., 2010). We showed ATP production was increased approximately 20% with cortisol treatment while reduced with RU 486 pretreatment (Fig. 5B). ATP upregulation generally leads to the deactivation of AMPK, which modulates the mTOR pathway. We showed that cortisol treatment decreased AMPK activity, but stimulated mTOR phosphorylation (Fig. 5C), which was remarkably reduced to the control level with xestospongin C or ruthenium red pretreatment (Fig. 5D). mTOR inhibits autophagy function by directly phosphorylating or inhibiting ULK, AMBRA1, or Atg14 (Galluzzi, Pietrocola et al., 2014). Our results also showed the decreased autophagy function representing Atg5 reduction, increased p62 level, and decreased LC3B formation (Fig. 5E). Generally, autophagy takes an important role in decreasing toxic Aβ. With impaired autophagy function due to cortisol, the decreased autophagosome formation around Aβ is significantly decreased and recovered by rapamycin pretreatment (Fig. 5F). Autophagy is also closely related to regulating protein level especially in the microtubule-associated proteins as cytoskeleton assembly plays a critical role in cell growth (He, Ding et al., 2016). Unlike mRNA expression of microtubule-associated proteins that remain unchanged (Fig EV. 3D), the expression of SCG10 was increased with cortisol (Fig. 5G) and downregulated with rapamycin pretreatment (Fig. 5H) where cortisol had no effect on stathmin-1. We therefore speculated that selective ubiquitination of SCG10 generally occurred, but was inhibited with cortisol. We showed that the ubiquitination of SCG10 was reduced with cortisol, which was increased to the control level upon rapamycin pretreatment (Fig. 5I). Co-localization of LC3 and SCG10 was also increased to the control level due to pretreatment of rapamycin (Fig. 5J). Eventually, increased SCG10-tubulin binding was observed due to upregulation of SCG10 level (Fig. 5K). Cortisol-regulated SCG10 was dependent on the ubiquitination-proteasome system as MG 132 pretreatment decreased the SCG10 level whereas cycloheximide had no effect on SCG10 expression (Fig EV. 3E).

**Figure 5.**
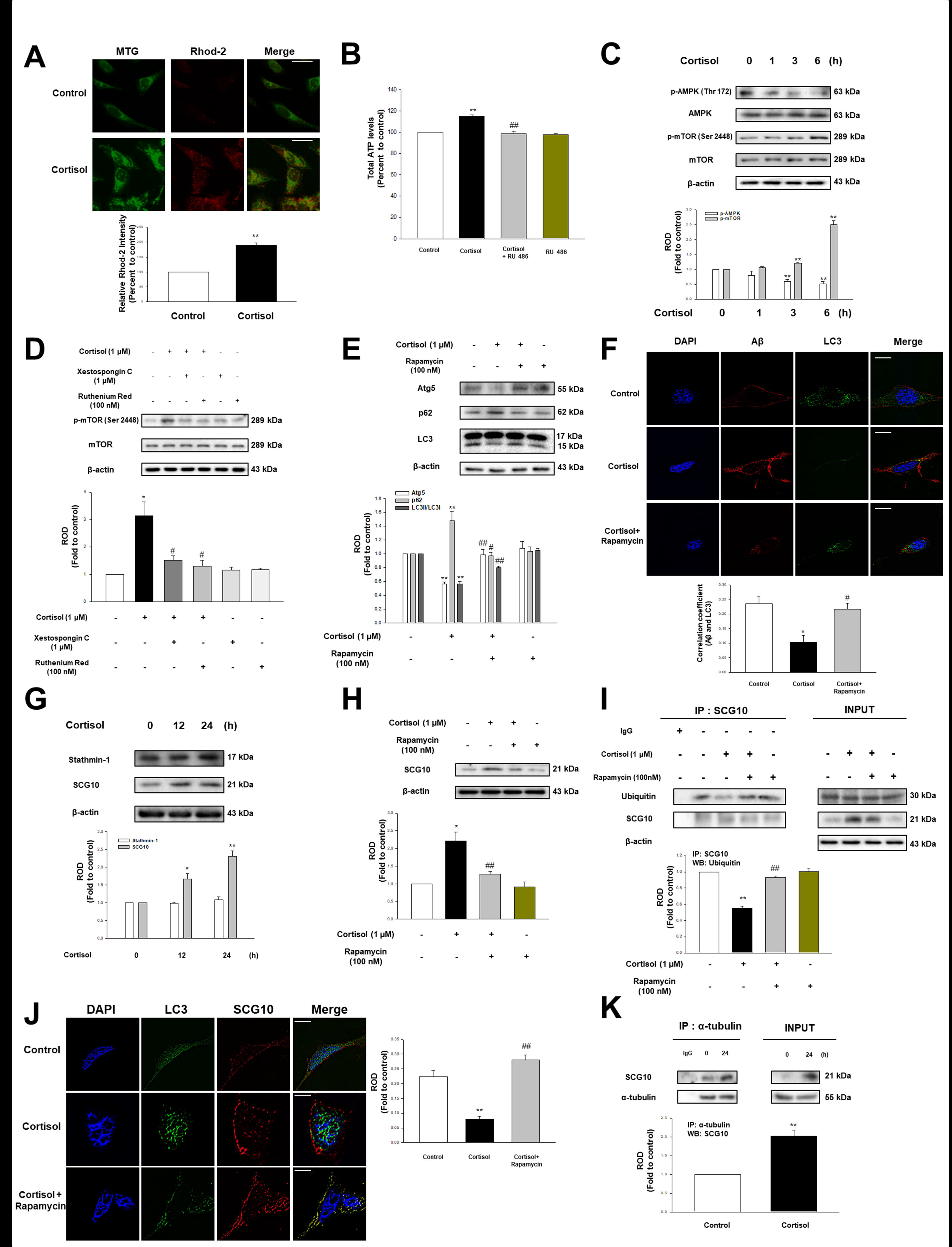
Cortisol inhibited autophagy via activation of mTOR. (**A**) The cells were treated with cortisol (1 μM) for 3 h, and stained with rhod-2 (3 μM) for 1 h to detect mitochondrial Ca^2+^. After incubation, mitotracker green (MTG, 300 nM) was also stained to visualize mitochondria. The intensity of both rhod-2 (red) and MTG (green) was measured with Eclipse Ts2™ fluorescence microscopy. Data are reported as a mean ± SE of five independent experiments. ** indicates *p<0.01* versus control. Scale bars represent 100 μm (magnification, × 400). (**B**) The cells were treated with RU 486 (1 μM) for 30 min before cortisol (1 μM) for 6 h, and then reacted with ATP luciferase reagent. The ATP levels were detected with luminometer. Data are reported as a mean ± SE of six independent experiments. ** indicates *p<0.01* versus control and ^##^ indicates *p<0.01* versus cortisol alone. (**C**) Time responses (0 - 6 h) of cortisol (1 μM) in phosphorylation of AMPK at Thr 172 and mTOR at Ser 2448 were shown. Data are reported as a mean ± SE of four independent experiments. ** indicates *p<0.01* versus control. (**D**) The cells were treated with xestospongin C (1 μM) for 2 h or ruthenium red (100 nM) for 30 min before cortisol (1 μM) for 6 h. p-mTOR (Ser 2448), mTOR, and β-actin were detected in western blotting results. Data are reported as a mean ± SE of four independent experiments. * indicates *p<0.05* versus control and ^#^ indicates *p<0.05* versus cortisol alone. (**E**) The cells were pretreated with rapamycin (100 nM) for 30 min before cortisol (1 μM) for 24 h. Atg5, p62, LC3, and β-actin were detected with western blot. Data are reported as a mean ± SE of five independent experiments. ** indicates *p<0.01* versus control. ^#^, ^##^ indicates *p<0.05, p<0.01* versus cortisol, respectively. (**F**) The cells were pretreated with rapamycin (100 nM) for 30 min before cortisol (1 μM) for 24 h. Co-localization of Aβ (red) and LC3 (green) was visualized with SRRF imaging system. DAPI was used for nuclear counterstaining (blue). Data are reported as a mean ± SE of five independent experiments. * indicates *p<0.05* versus control and ^#^ indicates *p<0.05* versus cortisol alone. Scale bars represent 20 μm (magnification, × 1,000). (**G**) Time responses (0 - 24 h) of cortisol (1 μM) in stathmin-1 and SCG10 expressions were shown. Data are reported as a mean ± SE of five independent experiments. *, ** indicates *p<0.05, p<0.01* versus control, respectively. (**H**) The cells were pretreated with rapamycin (100 nM) for 30 min before cortisol (1 μM) for 24 h. SCG10 and β-actin were detected with western blot. Data are reported as a mean ± SE of four independent experiments. * indicates *p<0.05* versus control and ^##^ indicates *p<0.01* versus cortisol. (**I**) The cells were pretreated with rapamycin (100 nM) for 30 min before cortisol (1 μM) for 24 h. SCG10 was co-immunoprecipitated with an anti-ubiquitin antibody (the left side). Expression of ubiquitin, SCG10, and β-actin in total cell lysates is shown in the right side. Data are reported as a mean ± SE of four independent experiments. ** indicates *p<0.01* versus control and ^##^ indicates *p<0.01* versus cortisol alone. (**J**) The cells were pretreated with rapamycin (100 nM) for 30 min before cortisol (1 μM) for 24 h. Co-localization of LC3 (green) and SCG10 (red) was visualized with SRRF imaging system. DAPI was used for nuclear counterstaining (blue). Data are reported as a mean ± SE of five independent experiments. ** indicates *p<0.01* versus control and ^##^ indicates *p<0.01* versus cortisol alone. Scale bars represent 20 μm (magnification, × 1,000). (**K**) The cells were treated with cortisol (1 μM) for 24 h and lysed. α-tubulin was co-immunoprecipitated with an anti-SCG10 antibody (the left side). Expression of SCG10 and α-tubulin in total cell lysates is shown in the right side. Data are reported as a mean ± SE of four independent experiments. ** indicates *p<0.01* versus control.

### Glucocorticoid triggers microtubule dysfunction and kinesin-1 detachment leading to memory deficits

Microtubule destabilization occurs when binding with Aβ or SCG10 through activating/deactivating enzymes that regulate the PTM of α-tubulin. Therefore, we showed that the decreased A/T ratio of α-tubulins in cells upon cortisol was restored with pretreatment of DAPT and rapamycin (Fig. 6A - 6B). Microtubule destabilization has been generally known to reduce the binding of kinesin-1, the motor protein that transports protein from cytosol close to the nucleus toward cellular terminus. We showed decreased binding of kinesin-1 to α-tubulin with cortisol (Fig. 6C - 6D). Our animal studies also showed decreased A/T ratio in the hippocampus of mice exposed to corticosterone. On the other hand, DAPT and rapamycin pretreatment restored A/T ratio (Fig. 6E). Moreover, kinesin-1 binding to α-tubulin was also recovered with DAPT and rapamycin pretreatment (Fig. 6F). As microtubule transports the important molecules into the necessary parts, we found that the memory-related receptors, AMPAR1/2 were not closely interacting with α-tubulin upon cortisol treatment (Fig. 6G – 6H). Mitochondria transport towards the cell terminus was also decreased by cortisol (Fig. 6I). Subsequently, the cell viability was reduced due to cortisol treatment, which was reversed by paclitaxel (microtubule stabilizer) pretreatment (Fig. 6J).

**Figure 6.**
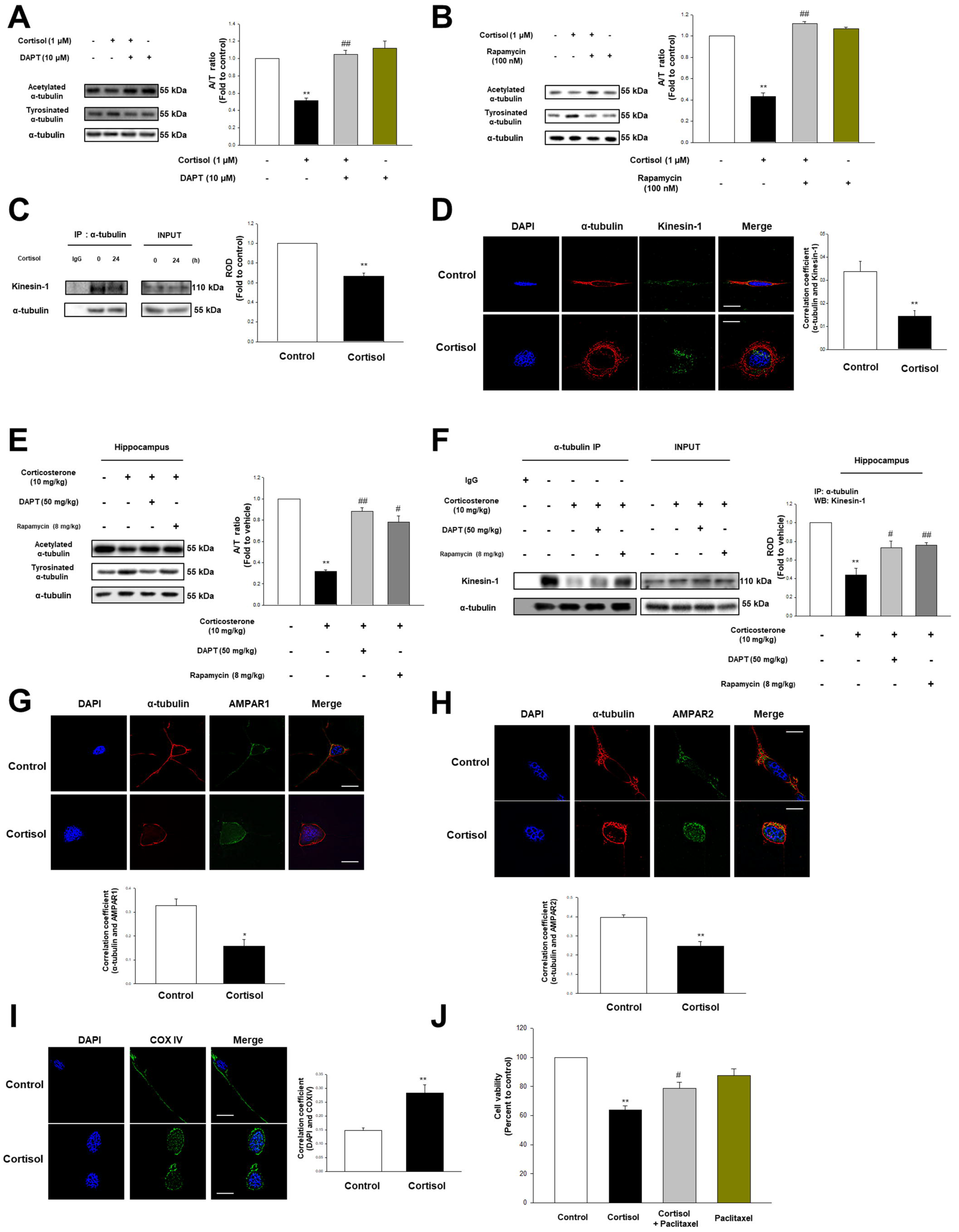
Glucocorticoid promoted microtubule destabilization and following transport impairment. (**A**) The cells were pretreated with DAPT (10 μM) for 30 min before cortisol (1 μM) for 48 h. Acetylated α-tubulin, tyrosinated α-tubulin, and α-tubulin were detected with western blot. Data are reported as a mean ± SE of four independent experiments. ** indicates *p<0.01* versus control and ^##^ indicates *p<0.01* versus cortisol alone. (**B**) The cells were pretreated with rapamycin (100 nM) for 30 min before cortisol (1 μM) for 48 h. Acetylated α-tubulin, tyrosinated α-tubulin, and α-tubulin were detected with western blot. Data are reported as a mean ± SE of four independent experiments. ** indicates *p<0.01* versus control and ^##^ indicates *p<0.01* versus cortisol alone. (**C**) The cells were treated with cortisol (1 μM) for 24 h. α-tubulin was co-immunoprecipitated with an anti-kinesin-1 antibody (the left side). Expression of kinesin-1 and α-tubulin in total cell lysates is shown in the right side. Data are reported as a mean ± SE of four independent experiments. ** indicates *p<0.01* versus control. (**D**) The cells were incubated with cortisol (1 μM) for 24 h. Co-localization of α-tubulin (red) and kinesin-1 (green) was visualized with SRRF imaging system. DAPI was used for nuclear counterstaining (blue). Data are reported as a mean ± SE of five independent experiments. ** indicates *p<0.01* versus control. Scale bars represent 20 μm (magnification, × 1,000). (**E**) The hippocampus of mice exposed to DAPT (50 mg/kg) for 2 h or rapamycin (8 mg/kg) for 2 days before corticosterone (10 mg/kg) for 24 h was collected and lysed. Acetylated α-tubulin, tyrosinated α-tubulin, and α-tubulin were detected by western blot. Data are reported as a mean ± SE of five independent experiments. ** indicates *p<0.01* versus vehicle. ^#^, ^##^ indicates *p<0.05, p<0.01* versus corticosterone alone, respectively. (**F**) The hippocampus of mice exposed to DAPT (50 mg/kg) for 2 h or rapamycin (8 mg/kg) for 2 days before corticosterone (10 mg/kg) for 24 h was collected and lysed. α-tubulin was co-immunoprecipitated with an anti-kinesin-1 antibody (the left side). Expression of kinesin-1 and α-tubulin in total cell lysates is shown in the right side. Data are reported as a mean ± SE of four independent experiments. ** indicates *p<0.01* versus vehicle. ^#^, ^##^ indicates *p<0.05, p<0.01* versus corticosterone alone, respectively. (**G**) The cells were incubated with cortisol (1 μM) for 48 h. Co-localization of α-tubulin (red) and AMPAR1 (green) was visualized with SRRF imaging system. DAPI was used for nuclear counterstaining (blue). Data are reported as a mean ± SE of five independent experiments. * indicates *p<0.05* versus control. Scale bars represent 20 μm (magnification, × 1,000). (**H**) The cells were incubated with cortisol (1 μM) for 48 h. Co-localization of α-tubulin (red) and AMPAR2 (green) was visualized with SRRF imaging system. DAPI was used for nuclear counterstaining (blue). Data are reported as a mean ± SE of five independent experiments. ** indicates *p<0.01* versus control. Scale bars represent 20 μm (magnification, × 1,000). (**I**) The cells treated with cortisol (1 μM) for 48 h were immunostained with DAPI (blue) and COX IV (green). Data are reported as a mean ± SE of five independent experiments. ** indicates *p<0.01* versus control. Scale bars represent 20 μm (magnification, × 1,000). (**J**) The cells were pre-incubated with paclitaxel (10 μM) for 30 min before cortisol (1 μM) for 48 h. After treatment, water soluble tetrazolium salt (WST-1) assay was performed to measure cell viability. Data are reported as a mean ± SE of six independent experiments. ** indicates *p<0.01* versus control and ^#^ indicates *p<0.05* versus cortisol alone.

We investigated whether microtubule stabilization reversed corticosterone treatment effect that induced memory impairment and following neurodegeneration *in vivo*. Corticosterone-treated mice group showed decreased translocation of AMPAR and TOMM20 into the synapse, which was recovered by paclitaxel injections (Fig. 7A). Similarly, reduced co-localization between synaptic marker (post synaptic density 95-PSD95) and TOMM20 or AMPAR1/2 was examined in hippocampal tissue of mice treated with corticosterone, which was increased with paclitaxel treatment (Fig. 7B - 7C). The TUNEL assay results showed that neuronal apoptosis was increased with corticosterone, which was reduced by paclitaxel treatment (Fig. 7D). In Y-maze test, mice injected with corticosterone showed memory impairment whereas the mice with pretreatment of paclitaxel showed recovered memory function (Fig. 7E). Overall, our data supported that microtubule dysfunction led to the failure of AMPAR or mitochondria trafficking into cell terminus, which promoted memory impairment.

**Figure 7.**
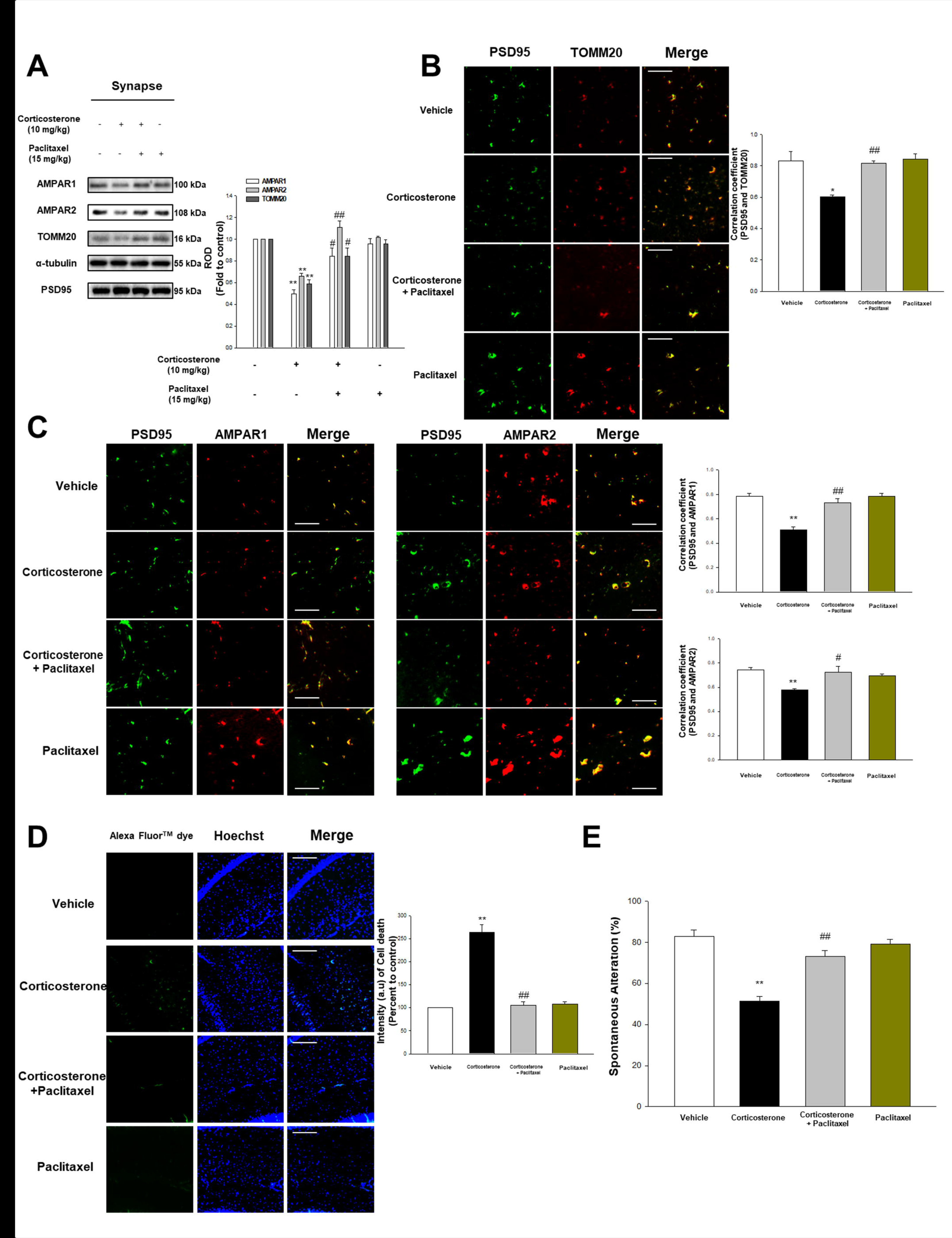
Corticosterone-induced memory impairment was attenuated by paclitaxel. (**A**) The synaptosome of hippocampus from mice exposed to vehicle, corticosterone (10 mg/kg), corticosterone with paclitaxel (15 mg/kg), and paclitaxel alone was isolated, and then AMPAR1, AMPAR2, TOMM20, α-tubulin, and PSD95 were detected. PSD95 was used as a loading control of synaptosome. Data are reported as a mean ± SE of six independent experiments. ** indicates *p<0.01* versus vehicle. ^#^, ^##^ indicates *p<0.05, p<0.01* versus corticosterone treatment group, respectively. (**B**) The mice were exposed to vehicle, corticosterone (10 mg/kg), corticosterone with paclitaxel (15 mg/kg), or paclitaxel. Slide samples for IHC were immunostained with PSD95 (green) and TOMM20 (red). Data are reported as a mean ± SE of five independent experiments. * indicates *p<0.05* versus vehicle and ^##^ indicates *p<0.01* versus corticosterone alone. Scale bars, 200 μm (magnification, × 200). (**C**) Slide samples for IHC were immunostained with PSD95 (green) and AMPAR1/2 (red). Data are reported as a mean ± SE of five independent experiments. ** indicates *p<0.01* versus vehicle. ^#^, ^##^ indicates *p<0.05, p<0.01* versus corticosterone alone, respectively. Scale bars, 200 μm (magnification, × 200). (**D**) TUNEL assay was performed using slide samples of hippocampus from mice with vehicle, corticosterone (10 mg/kg), corticosterone with paclitaxel (15 mg/kg), or paclitaxel. The intensity of green fluorescence indicates the amount of neuronal cell death. Data are reported as a mean ± SE of five independent experiments. ** indicates *p<0.01* versus vehicle and ^##^ indicates *p<0.01* versus corticosterone alone. Scale bars, 200 μm (magnification, × 200). (**E**) The mice exposed to vehicle, corticosterone (10 mg/kg), corticosterone with paclitaxel (15 mg/kg), or paclitaxel were subjected to Y-maze test to evaluate memory function. Data are reported as a mean ± SE of six independent experiments. ** indicates *p<0.01* versus vehicle and ^##^ indicates *p<0.01* versus corticosterone alone.

## Discussion

Our study not only showed the glucocorticoid-mediated changes in microtubule dynamics through increased ER-mitochondria coupling followed by Aβ accumulation and SCG10 upregulation, but also demonstrated how microtubule dysfunction affects memory formation in both the animal model and SH-SY5Y cells. We first demonstrated that microtubule destabilization in the hippocampus of mice with corticosterone eventually induced memory deficits through failed AMPAR or mitochondria transport. There are considerable research demonstrating how glucocorticoid triggers microtubule dysfunction via amyloidosis and tau hyperphosphorylation by genomic pathway (Green et al., 2006) or membrane GR-dependent CREB pathway (Choi, Lee et al., 2017). However, current research elucidated that the mechanisms of mitochondrial GR on the microtubule dysfunction. Mitochondrial GR can directly affect the mitochondrial function like transcription of OXPHOS genes, ATP synthesis, and Ca^2+^ reuptake (Hunter, Seligsohn et al., 2016, Psarra & Sekeris, 2011, Yao, Irwin et al., 2009). These changes in mitochondrial function alter microtubule dynamics since assembly or PTM of tubulin is mediated by mitochondrial Ca^2+^, which in turn affects ATP synthesis (Du, Wang et al., 2009b, McEwen, Bowles et al., 2015). There are many debates considering whether or not cortisol induced the translocation of GR into mitochondria. Chronic cortisol treatment triggered the reductions in GR trafficking into mitochondria due to the changes in GR expressions or PTM (Du et al., 2009b). However, with short term treatment representing an acute stress state, cortisol usually stimulated GR movement into mitochondria, which is the same as our condition. In particular, GR designated toward mitochondria has been widely accepted for binding to Bcl-2 and moving to ER or mitochondria for regulating mitochondrial calcium capacity, but the detailed mechanism has not been implicated (Picard & McEwen, 2014). In this paper, we demonstrate how the GR-Bcl-2 complex increased the mitochondrial calcium sequestration via upregulating ER-mitochondria connectivity. Normally, about twenty percent of mitochondria are closely related to MAM but increased ER-mitochondria communication has been extensively observed in Alzheimer’s disease models, indicating that Alzheimer’s disease is deeply associated with the MAM function (Pinho, Teixeira et al., 2014, Schon & Area-Gomez, 2013). Interestingly, our works showed that ER-mitochondria contacts formed by GR-Bcl-2 complex triggered IP3R-VDAC1 bridging where both GR and Bcl-2 attach. Bcl-2 is previously described as binding to IP3R and mediating Ca^2+^ transfer from ER into mitochondria (Cárdenas & Foskett, 2012, Hedskog et al., 2013, Krugers, Hoogenraad et al., 2010). Surprisingly, we also suggested GR interacts with IP3R, which was not observed in previous research. Indeed, we could assume that GR-Bcl-2 complex binds to the MAM region serving as scaffold to help ER-mitochondria coupling but the exact mechanism needs further investigation. Several tethering proteins function to increase interaction between MAM and mitochondria such as PACS2-mfn2 and VAPB-PTPIP51 (Paillusson, Stoica et al., 2016). In our work, cortisol formed mfn2-PACS2 protein complex, one of the major tethering protein complexes stabilizing the ER-mitochondria network that was decreased with knockdown of *bcl-2* (Area-Gomez, De Groof et al., 2009, Hedskog et al., 2013). We also confirmed that inhibiting ER-mitochondria coupling restored A/T ratio suggesting that glucocorticoid-induced GR translocation into mitochondria is highly associated with microtubule destabilization through integration of ER and mitochondria. We suggest this as the major hallmark of stress-related Alzheimer’s disease.

Recent studies have concentrated on elucidating new therapeutic approach to Alzheimer’s disease through an in-depth inspection on the downstream effect of ER-mitochondria interaction since it is pathogenic change of Alzheimer’s disease and precedes the accumulation of plaques or tangles (Pinho et al., 2014). Firstly, we demonstrated that increased ER-mitochondria interaction by cortisol can induce Alzheimer’s disease through γ-secretase-dependent Aβ production. The role of γ-secretase in glucocorticoid-induced amyloidosis has not been studied in depth since there were no changes in its expression. Thus, there have been few investigations into the stimulating effect of glucocorticoid on γ-secretase in T cell lymphoma treatment (Real, Tosello et al., 2009). However, γ-secretase itself can trigger amyloidosis through cleaving C99 into Aβ when localized in MAM by increasing its activities. Our works also exhibited that γ-secretase localizes ER-mitochondria contact sites to increase Aβ formation. Previous research also suggested that cleavage of C99 with γ-secretase at MAM resulted in the Aβ transport in mitochondria through TOM complex and subsequent extracellular Aβ accumulation (Pera, Larrea et al., 2017, Schreiner, Hedskog et al., 2015). Consistent with these reports, we demonstrated that cortisol induced co-localization of γ-secretase or C99 at MAM, and also showed accumulated Aβ in mitochondria via upregulated γ-secretase activity due to increased ER-mitochondria interaction. Inhibiting GR-Bcl-2 complex to avoid ER-mitochondria interaction or downregulating γ-secretase activity also dramatically decreased C99 localization and Aβ production at MAM which further suggests the novel therapeutic approach to AD. Secondly, ER-mitochondria interaction via IP3R-VDAC1 bridging also increases mitochondrial Ca^2+^ which finally leads to autophagy dysfunction (Cárdenas & Foskett, 2012, Cárdenas et al., 2010, Gomez-Suaga, Paillusson et al., 2017). Ca^2+^ overload in mitochondria conversely leads to increased ATP synthesis via stimulating OXPHOS enzymes (Pera et al., 2017). Although some previous reports suggested that chronic glucocorticoid treatment reduced ATP synthesis due to prolonged mitochondria damage and different tendency of regulating OXPHOS genes, many studies showed glucocorticoid promotes ATP synthesis by increasing its calcium storing capacity and GR binding to the promoters of OXPHOS genes in mitochondrial DNA (Hunter et al., 2016, Psarra & Sekeris, 2011). With ATP level upregulation, the deactivation of AMPK either phosphorylates TSC2 or raptor to finally inhibit the autophagy process (Galluzzi et al., 2014). Several studies showed that glucocorticoid elevated autophagy-related genes stimulating autophagy process in lymphocytes and osteoblasts (Molitoris, McColl et al., 2011, Shen, Ren et al., 2018). However, the glucocorticoid dosage to treat such inflammatory diseases is different from the stress-induced glucocorticoid level since it is designed to efficiently induce autophagic cell death. In contrast, our results showed that the deactivation of AMPK triggered phosphorylating its substrate, mTOR, leading to defective autophagy, similar to glucocorticoid effect on mTOR signaling in the previous report (Wang, Wang et al., 2017). Subsequent autophagy dysfunction failed to evoke Aβ clearance in SH-SY5Y cells, which agrees with the observations including altered mTOR signaling and subsequent autophagy defects in Alzheimer’s disease (Caccamo, Magrì et al., 2013). Furthermore, we also showed that glucocorticoid inhibited selective autophagy towards the SCG10. Selective autophagy via ubiquitination is important to avoid the accumulation of redundant proteins to avoid cellular damage (Shaid, Brandts et al., 2013, Song, Yi et al., 2016). In particular, selective autophagy serve as crucial regulator of microtubules responsible for migration, development, and differentiation (Grenningloh, Soehrman et al., 2004, Nunan, Shearman et al., 2001). There are many proteins subject to selective autophagy inducing axonal degeneration such as α-synuclein, huntingtin proteins, stathmin, and neurofilament. Namely, if defective autophagy clearance occurs, the microtubule dysfunction follows. Previous reports suggest that autophagy induction stabilize neuronal microtubule via decreasing SCG10, indicating that autophagy plays a pivotal role in regulating SCG10 level (He et al., 2016, Shin, Geisler et al., 2014). Our results also indicated that ubiquitination and selective autophagy of SCG10 among stathmin family was reduced by cortisol whereas inhibition of mTOR triggered autophagy induction to eventually reduce SCG10 level.

Aβ or SCG10-dependent changes in microtubule dynamics were already reported to be involved in memory formation, but the exact mechanism of how glucocorticoid induces memory deficits via microtubule dysfunction is far from clear. Mounting evidence also suggest that multiple bindings between kinesin-1 and α-tubulin moves necessary components further than single binding, indicating that the kinesin-1 detachment from α-tubulin can dramatically fail to transfer cargos into cell terminus (Vershinin, Carter et al., 2007). Thus, microtubule destabilization due to increased tyrosination of α-tubulin triggers reduction in the binding of kinesin-1 motor protein to α-tubulin (Uchida et al., 2014). Accordingly, we exhibited that decreasing Aβ or SCG10 level restored the A/T ratio and kinesin-1 binding to α-tubulin. Aβ is well known to induce various mechanisms to trigger microtubule instability. Aβ itself functions as microtubule depolymerizer, activator of kinases which trigger tau hyperphosphorylation, or receptor-mediated destabilizer to change microtubule dynamics (Baas & Ahmad, 2013). Furthermore, stathmin-dependent changes in microtubule stability influence memory formation. Some reports showed that unstable microtubule is necessary for memory consolidation and loss of stathmin level evokes memory loss (Uchida et al., 2014). However, this report focused on the phosphorylation state of stathmin which inactivates the depolymerizing activity of stathmin. On the other hand, previous research reported that excessive SCG10-tubulin binding significantly affects the microtubule stability, leading to decreased transport at the synaptic sites (Uchida et al., 2014). Given the observations, we speculated that the cells are likely to fail transporting the necessary components towards the cell periphery. Microtubule transports the memory-related receptors and mitochondria to establish the memory consolidation and synaptic strength (Kaganovsky & Wang, 2016). For example, the stathmin family regulates binding between AMPAR2 and KIF5 into synaptic sites maintaining memory function (Dent, 2017). Localization of AMPARs in the synaptic parts plays a major role in establishing memory and maintaining hippocampal long-term potentiation (Brown, Tran et al., 2005, Krugers et al., 2010, Lu, Man et al., 2001). However, the relationship between glucocorticoid and AMPAR has not yet been studied. The *in vitro* and *in vivo* studies demonstrated that the AMPAR transport and mitochondria into cellular extremities and synapse was significantly reduced upon glucocorticoid treatment. Furthermore, these works agree with the previous report demonstrating that excessive glucocorticoid triggers the synaptic dysfunction and hippocampal apoptosis with increased caspases (Sotiropoulos, Silva et al., 2015), while physiological levels of glucocorticoid facilitates the memory consolidation. We used microtubule-stabilizing drug for helping microtubule become acetylated and providing benefits to Alzheimer’s disease (Baas & Ahmad, 2013). Using paclitaxel that stimulates the recovery of the AMPAR and mitochondria levels, memory formation was restored to normal functions and the neuronal cell death was reduced, indicating that microtubule dynamics play an important role in memory function. Therefore, our data demonstrate that glucocorticoid impairs the localization of memory-related receptors and mitochondria in the synapse by Aβ or SCG10-mediated changes in microtubule stability.

In conclusion, the results of this study showed that glucocorticoid impairs microtubule function by reorganization the ER-mitochondria interaction which leads to Aβ formation at MAM and selective autophagosome formation of SCG10. Furthermore, we also demonstrated the underlying mechanisms that microtubule destabilization by glucocorticoid finally decreases memory formation and induces neurodegeneration via inhibiting the transport of AMPAR and mitochondria into the cell terminus. Thus, our approach to investigate the novel signaling pathways of glucocorticoid on Alzheimer’s disease via microtubule dysfunction in the animal model and neuroblastoma cells can provide potential therapeutic targets that control the microtubule stability.

## Materials and Methods

### Materials

Cells from the human neuroblastoma cell line SH-SY5Y were obtained by Korean Cell Line Bank (Seoul, Korea). Fetal bovine serum (FBS) and serum replacement (SR) were purchased from Hyclone (Logan, UT, USA) and Gibco (Grand Island, NY, USA), respectively. The antibodies of mfn2, presenilin1 (PSEN1), VDAC1, stathmin-1, SCG10, Bcl-2, p-mTOR (Ser 2448), p-AMPK (Thr 172), and β-actin were acquired from Santa Cruz Biotechnology (Santa Cruz, CA, USA). The antibodies of TOMM20, IP3R, Aβ, and the product of xestospongin C were purchased from Abcam (Cambridge, England). Horseradish peroxidase (HRP)-conjugated goat anti-rabbit IgG was obtained from Jackson Immunoresearch (West Groove, PA, USA). The antibodies of AMPAR1, AMPAR2, C99, and PSD95 were purchased from EMD Millipore (Billerica, MA, USA). The antibodies of AMPK, mTOR, Atg5, and GR were purchased from Cell Signaling Technology, Inc. (Danvers, MA, USA). The antibodies of LC3, p62, and kinesin-1 were purchased from Novus Biologicals (Littleton, CO, USA). The antibodies of ubiquitin, tyrosinated α-tubulin, and PSD95 were obtained from Thermo Fisher (Rockford, IL, USA). The antibodies of COX IV, Hsp90, Hsp70, PACS2, PTPIP51, and the chemical cortisol-BSA were obtained from Cusabio (Wuhan, Hubei, China). Cortisol, corticosterone, BSA, RU 486, paclitaxel, DAPI, DAPT, actinomycin D, rapamycin, cycloheximide, MG132, Hoechst, ruthenium red, and the antibodies of acetylated α-tubulin, α-tubulin were purchased from Sigma Chemical Company (St. Louis, MO, USA). VER 155008 was obtained from Calbiochem (Merk Millipore, Darmstadt, Germany).

### Cell Culture

The SH-SY5Y cells were cultured in high glucose Dulbecco’s Modified Eagle Medium (DMEM, Hyclone) containing 10% FBS and 1% antibiotic-antimycotic mixture. Cells were cultured in 60, 100 mm diameter culture dishes, or a 96-well plate in an incubator kept at 37 °C with 5% CO_2_. Cells were incubated for 72 h and then the medium was replaced with serum-free DMEM containing 1% SR and 1% antibiotic-antimycotic solution for 24 h.

### Experimental Design

Male ICR mice exposed to corticosterone mimic the stress-induced mouse model since corticosterone is primarily responsive steroid hormone to stress. The hippocampus of mice was mainly used for evaluating glucocorticoid effect on microtubule dynamics since the hippocampus has the most abundant GRs among the brain.

### Animal preparation and drug treatment

Male ICR mice aged 7 weeks were used, in compliance and approval with the Institutional Animal Care and Use Committee of Seoul National University (SNU-171017-9). Animals were housed 6 per cage under standard environmental conditions (22 °C relative humidity 70%; 12 h light: dark cycle; *ad libitum* access to food and drinking solution). Corticosterone (10 mg/kg) was dissolved in the solution containing 50% propylene glycol in PBS and injected intraperitoneally (Choi et al., 2017). Vehicle-treated mice were similarly injected with the solution containing propylene and PBS. DAPT (50 mg/kg) and rapamycin (8 mg/kg) were dissolved in the solution containing 1% DMSO and 99% corn oil (Sigma Chemical Company). The dosage and treatment period of DAPT and rapamycin is modified from previous reports (Bittner, Fuhrmann et al., 2009, Malagelada, Jin et al., 2010). The vehicle solution includes the 1% DMSO and 50% propylene glycol in PBS. Paclitaxel (15 mg/kg) was dissolved in the solution containing 50% β-cyclodextrin and treated 3 h before the corticosterone treatment following the modified method and dosage of previous report (Fellner, Bauer et al., 2002). Vehicle-treated mice were similarly injected with the solution containing propylene glycol and β-cyclodextrin. Mice were monitored twice a day during all experiments.

### Sample Size

Total seventy two of 7-week-old male ICR mice were used for the *in vivo* study. Six mice were utilized for each group throughout the study. Applying size of samples (minimum of n = 3) can be acceptable if very low *p* values are observed rather than the large size of N including interfering results (Vaux, 2012). Therefore, we set the minimum of n = 4 (western blotting, IHC) and n = 6 (behavior test) to gain statistical powers according to the previous published article of brain (Olmos-Alonso, Schetters et al., 2016).

### Randomization and blinding

The experiments were designed in compliance with the ARRIVE guidelines. Allocations of animals were randomly done to minimize the effects of subjective bias. No exclusion of data obtained from samples was done.

### Y-Maze spontaneous alteration test

Y-maze spontaneous alteration test is based on the innate willingness of rodents to differently explore new environments. This behavior test is frequently used for quantifying the cognitive deficits of the animals. Rodents usually prefer to challenge a new arm of the Y-maze rather than returning back to the one which was previously visited. Before the test, the animals were placed in the home cage at the testing room for 3 h to minimize the effects of stress on behavior. The mice were placed in the Y-shaped maze purchased from Sam-Jung Company (Seoul, Korea). The mice were allowed to explore the open field for 10 min, and at the same time, the number of arm entries and triads were recorded to calculate the percentage of alteration. Only an entry when all four limbs were within the arm was counted. The alteration amount represents the number of alterations which was divided by total triads (total entries – 2). When the animals show higher alteration percentage, the animals tend to maintain the memory function.

### Immunohistochemistry (IHC)

Mice underwent deep anesthesia with zoletil (50 mg/kg) and were perfused transcardially with calcium-free Tyrode’s solution followed by a fixative including 4% paraformaldehyde in 0.1 M phosphatebuffer (pH 7.4). The removed brain was post-fixed for 2 h, and subsequently placed in 30% sucrose in PBS for 24 h at 4 °C. Serial transverse sections (40 μm) were performed using a cryostat (Leica Biosystems, Nussloch, Germany). The brain tissues containing hippocampus were fixed with 4% paraformaldehyde and then pre-blocked with 5% normal goat serum (Sigma Chemical Company) containing 0.1% Triton X-100 in PBS at room temperature for 1 h. Samples were incubated with primary antibodies overnight at 4 °C, followed by secondary antibodies for 2 h at room temperature. Completed samples were visualized by using Eclipse Ts2™ fluorescence microscopy (Nikon, Tokyo, Japan). The fluorescent intensity analysis was undertaken using Fiji software.

### Synaptic protein extraction

Synaptosome of hippocampus was extracted using Syn-Per synaptic protein extraction reagent (Thermo Fisher). Hippocampus was homogenized with Dounce grinder with 20 slow strokes. The homogenates underwent centrifugation at 1,200 × g for 10 min at 4 °C. After discarding the pellet, the supernatant was centrifuged at 15,000 × g for 20 min at 4 °C. The supernatant part is cytosolic fraction and the pellet (synaptosome) was dissolved with the extraction reagent.

### TUNEL assay

The cryo-sectioned brain tissue underwent the TUNEL assay using Click-iT^TM^ TUNEL Alexa Fluor^TM^ 488 Imaging Assay (Thermo Fisher) to evaluate the apoptosis of hippocampal neuronal cells. The samples were fixed with 4% paraformaldehyde for 15 min and then placed with 0.25% triton X-100 in PBS for 20 min at room temperature. The processes of TdT incorporation of EdUTP into dsDNA strand breaks and incubation with fluorescent dye detecting the EdUTP were conducted according to the manufacturer’s instructions. Images were obtained by Eclipse Ts2™ fluorescence microscopy. The fluorescent intensity analysis was performed using Fiji software.

### Reverse transcription-polymerase chain reaction (RT-PCR) and real-time PCR

The SH-SY5Y cells were treated for 12 h with cortisol, and the total RNA was extracted using MiniBEST Universal RNA Extraction Kit (TaKaRa, Otsu, Shinga, Japan). Reverse transcription was performed using 1 μg of RNA with a Maxime RT-PCR PreMix Kit (Intron Biotechnology, Seongnam, Korea) to obtain cDNA. Two microliters of cDNA was then amplified using Quanti NOVA SYBR Green PCR Kits (Qiagen, Hilden, Germany). Real-time quantification of RNA targets was performed in a Rotor-Gene 6000 real-time thermal cycling system (Corbett Research, NSW, Australia). The primers for microtubule associated proteins were purchased from the Bioneer Corporation (Daejeon, Korea). The reaction mixture (20 μl) contained 200 ng of total RNA, 0.5 mM of each primer, and appropriate amounts of enzymes and fluorescent dyes as recommended by the manufacturer. The real-time PCR was performed as follows: 15 min at 95 °C for DNA polymerase activation; 15 sec at 95 °C for denaturing; and 50 cycles of 15 sec at 94 °C, 30 sec at 54 °C, and 30 sec at 72 °C. Data were collected during the extension step, and analysis was performed using the software provided; melting curve analysis was performed to verify the specificity and identity of the PCR products. Normalization of gene expression levels was performed by using the *β-actin* gene as a control.

### Western blot analysis

Harvested tissues or cells were incubated with the appropriate buffer for 30 min on ice. The lysates were cleared by centrifugation (10,000 × g at 4 °C for 30 min) and the supernatant was collected. To evaluate the protein concentration, the bicinchoninic acid (BCA) assay kit (Bio-Rad, Hercules, CA, USA) was used. Equal amounts of sample proteins (1 - 5 μg) were prepared for 8-15% SDS-PAGE and then transferred to a polyvinylidene fluoride membrane. Subsequently, the membranes were blocked with 5% BSA or 5% skim milk (Gibco) in TBST solution for 30 min. Blocked membranes were incubated with primary antibody overnight at 4 °C. The membranes were then washed and incubated with the HRP-conjugated secondary antibody at room temperature for 2 h. The western blotting bands were visualized by using chemiluminescence solution (Bio-Rad) and the densitometry analysis for quantification was carried out by using Image J software.

### Small interfering RNA (siRNA) transfection

Cells were grown until 60% of the surface of the plate. Prior to cortisol treatment, siRNAs specific for *bcl-2* and *nontargeting* (*NT*) obtained from Bioneer Corporation (Daejeon, Korea), and Dharmacon (Lafayette, CO, USA), respectively, were transfected to cells for 24 h with turbofect transfection reagent (Thermo Fisher) according to the manufacturer’s instructions. The concentration of each transfected siRNA was 25 nM. *NT* siRNA was used as the negative control.

### Immunocytochemistry

Cells on a confocal dish (Thermo Fisher) were fixed with 80% acetone for 10 min. Then, cells were incubated with 5% normal goat serum in PBS and incubated with primary antibody for overnight in 4 °C. Next, the cells were incubated for 2 h at room temperature with Alexa fluor secondary antibody. Images were obtained by Eclipse Ts2™ fluorescence microscopy or super-resolution radial fluctuations (SRRF) imaging system [(Andor Technology, Belfast, UK), (Gustafsson, Culley et al., 2016)]. The fluorescent intensity analysis and co-localization analysis with Pearson’s correlation coefficient were acquired by Fiji software.

### Measurement of cellular ATP levels

Intracellular ATP concentration level of cells was measured using ATP Bioluminescent HSII kit (Roche, Basel, Switzerland) according to the manufacturer’s instructions. ATP levels were detected with luminometer (Victor3; Beckman Coulter, Fullerton, CA, USA) and ATP concentrations were normalized to total protein concentration.

### Co-immunoprecipitation

The magnetic bead conjugated with specific primary antibodies was immobilized according to the supplier’s instructions. The total lysates of cells (300 μg) was incubated with 10 μg of primary antibody for overnight at 4 °C. Magnetic beads were spun-down by magnet and then collected. The antibody-bound protein was acquired by incubation in elution buffer (Thermo Fisher).

### In situ proximal ligation assay (PLA)

Duolink™ in situ PLA was performed according to the manufacturer’s instructions (Sigma Chemical Company). After fixation and blocking at 37 °C, primary antibodies against rabbit anti-IP3R and mouse anti-VDAC1 were diluted in antibody diluent and then incubated overnight at 4 °C. The Duolink™ secondary antibodies against the particular primary antibodies were applied for 1 h at 37 °C. DNA was then ligated for 1 h at 37 °C and incubated with polymerase. Fluorescent images were visualized with Eclipse Ts2™ fluorescence microscopy or SRRF imaging system.

### Measurement of Aβ in culture medium

Aβ (1-42) level in culture medium was determined using a human Aβ (1-42) ELISA kit from Wako Pure Chemical Industries, Ltd (Chuo-Ku, Osaka, Japan). Following the manufacturer’s instructions, the value of OD_450nm_ was measured and quantified into Aβ concentration, according to the standard curve.

### Water soluble tetrazolium salt (WST-1) assay

WST-1 assay was used for determining the cell proliferation and viability *in vitro* model. After treatment, cells were incubated in 10 μl of EZ-Cytox^TM^ solution including WST-1 in 100 μl of medium for 30 min at 37 °C with 5% CO_2_. The absorbance of each sample using a microplate reader was measured at a wavelength of 450 nm.

### Statistical analysis

Results are expressed as mean value ± standard error of mean (SE) and analyzed with the sigma plot 10 software. All experiments were analyzed by ANOVA, and some experiments which needed to compare with 3 groups were examined by comparing the treatment means to the control using a Bonferroni-Dunn test. A result with a *p* value of < 0.05 was considered statistically significant.

## Acknowledgements

This research was supported by National R&D Program through the National Research Foundation of Korea (NRF) funded by the Ministry of Science, ICT & Future Planning (NRF-2013M3A9B4076541 and NRF-2017R1A2B2008661). The funders had no role in study design, data collection or analysis, the decision to publish, or manuscript preparation.

## Author Contributions

Gee Euhn Choi: Designed research, data analysis and interpretation, performed experiments, wrote the paper

Ji Young Oh: Data analysis and interpretation, performed experiments

Hyun Jik Lee: Data analysis and interpretation

Chang Woo Chae: Performed experiments

Jun Sung Kim: Performed experiments

Ho Jae Han: Designed research, wrote the paper

## Expanded view figure legends

**Figure EV 1. Microtubule dysfunction by cortisol is mediated by mitochondrial GR in SH-SY5Y cells.** (**A**) The cells were incubated with actinomycin D (500 ng/ml) for 30 min before cortisol treatment (1 μM) for 48 h. Acetylated α-tubulin, tyrosinated α-tubulin, and α-tubulin were detected. Data are reported as a mean ± SE of four independent experiments. ** indicates *p<0.01* versus control. (**B**) The cells were treated with cortisol-BSA (1 μM) for 48 h. Acetylated α-tubulin, tyrosinated α-tubulin, and α-tubulin were detected. n=4. (**C**) The cells were incubated with RU 486 (1 μM) for 30 min before cortisol treatment (1 μM) for 2 h. GR, α-tubulin, and COX IV were detected. Cytosolic and mitochondrial protein expressions were normalized by α-tubulin and COX IV, respectively, in western blotting results. Data are reported as a mean ± SE of four independent experiments. * indicates *p<0.05* versus control and ^#^ indicates *p<0.05* versus cortisol. (**D**) The cells were incubated with VER 155008 (10 μM) for 30 min before cortisol treatment (1 μM) for 48 h. Acetylated α-tubulin, tyrosinated α-tubulin, and α-tubulin were detected. Data are reported as a mean ± SE of four independent experiments. ** indicates *p<0.01* versus control and ^##^ indicates *p<0.01* versus cortisol.

**Figure EV 2. Increased ER-mitochondria contact by cortisol induced microtubule destabilization.** (**A**) Knockdown of *bcl-2* was done using siRNA transfection for 24 h and then cells were treated with cortisol (1 μM) during 2 h. VAPB was co-immunoprecipitated with an anti-PTPIP51 antibody (the left side). Expression of PTPIP51, VAPB, and β-actin in total cell lysates is shown in the right side. n=4. (**B**) Knockdown of *bcl-2* was done using siRNA transfection for 24 h and then cells were treated with cortisol (1 μM) during 48 h. Acetylated α-tubulin, tyrosinated α-tubulin, and α-tubulin were detected. Data are reported as a mean ± SE of four independent experiments. ** indicates *p<0.01* versus control and ^##^ indicates *p<0.01* versus cortisol. (**C**) The cells were incubated with RU 486 (1 μM) for 30 min before cortisol treatment (1 μM) for 48 h. Acetylated α-tubulin, tyrosinated α-tubulin, and α-tubulin were detected. Data are reported as a mean ± SE of four independent experiments. ** indicates *p<0.01* versus control and ^##^ indicates *p<0.01* versus cortisol.

**Figure EV 3. Mitochondria Ca^2+^ induced by ER-mitochondria connectivity increased SCG10 level.** (**A**) Knockdown of *bcl-2* was done using siRNA transfection for 24 h and then cells were treated with cortisol (1 μM) during 3 h. Then the cells were stained with rhod-2 (3 μM) for 1 h to detect mitochondrial Ca^2+^ by luminometer. Data are reported as a mean ± SE of six independent experiments. ** indicates *p<0.01* versus control and ^##^ indicates *p<0.01* versus cortisol. (**B**) The cells were treated with xestospongin C (1 μM) for 2 h before cortisol (1 μM) for 3 h. Then the cells were stained with rhod-2 (3 μM) for 1 h to detect mitochondrial Ca^2+^ by luminometer. Data are reported as a mean ± SE of six independent experiments. ** indicates *p<0.01* versus control and ^##^ indicates *p<0.01* versus cortisol. (**C**) The cells were treated with ruthenium red (100 nM) for 30 min before cortisol (1 μM) for 3 h. Then the cells were stained with rhod-2 (3 μM) for 1 h to detect mitochondrial Ca^2+^ by luminometer. Data are reported as a mean ± SE of six independent experiments. * indicates *p<0.05* versus control and ^#^ indicates *p<0.05* versus cortisol. (**D**) The cells were treated with cortisol (1 μM) during 12 h, and mRNA was extracted. Real time PCR was performed to measure mRNA expressions. β-actin was used for loading control. n=6. (**E**) The cells were treated with cycloheximide C (1 μM) or MG 132 (100 nM) for 30 min before cortisol (1 μM) for 24 h. SCG10 and β-actin were detected in western blotting results. Data are reported as a mean ± SE of four independent experiments. . ** indicates *p<0.01* versus control and ^##^ indicates *p<0.01* versus cortisol.

## References

Akner G, Wikstro A-C, Stro P-E, Stockman O, Wallin M (1995) Glucocorticoid receptor inhibits microtubule assembly in vitro. Molecular and cellular endocrinology 110: 49–54

Area-Gomez E, De Groof AJ, Boldogh I, Bird TD, Gibson GE, Koehler CM, Yu WH, Duff KE, Yaffe MP, Pon LA (2009) Presenilins are enriched in endoplasmic reticulum membranes associated with mitochondria. The American journal of pathology 175: 1810–1816

Baas PW, Ahmad FJ (2013) Beyond taxol: microtubule-based treatment of disease and injury of the nervous system. Brain 136: 2937–2951

Bezprozvanny I, Mattson MP (2008) Neuronal calcium mishandling and the pathogenesis of Alzheimer’s disease. Trends in neurosciences 31: 454–463

Bittner T, Fuhrmann M, Burgold S, Jung CK, Volbracht C, Steiner H, Mitteregger G, Kretzschmar HA, Haass C, Herms J (2009) γ-secretase inhibition reduces spine density in vivo via an amyloid precursor protein-dependent pathway. Journal of Neuroscience 29: 10405–10409

Brandt R, Bakota L (2017) Microtubule dynamics and the neurodegenerative triad of Alzheimer’s disease: The hidden connection. Journal of neurochemistry 143(4): 409–417

Brown TC, Tran IC, Backos DS, Esteban JA (2005) NMDA receptor-dependent activation of the small GTPase Rab5 drives the removal of synaptic AMPA receptors during hippocampal LTD. Neuron 45: 81–94

Caccamo A, Magrì A, Medina DX, Wisely EV, López◻Aranda MF, Silva AJ, Oddo S (2013) mTOR regulates tau phosphorylation and degradation: implications for Alzheimer’s disease and other tauopathies. Aging cell 12: 370–380

Cárdenas C, Foskett JK (2012) Mitochondrial Ca^2+^ signals in autophagy. Cell calcium 52: 44–51

Cárdenas C, Miller RA, Smith I, Bui T, Molgó J, Müller M, Vais H, Cheung K-H, Yang J, Parker I (2010) Essential regulation of cell bioenergetics by constitutive InsP3 receptor Ca^2+^ transfer to mitochondria. Cell 142: 270–283

Choi GE, Lee S-J, Lee HJ, Ko SH, Chae CW, Han HJ (2017) Membrane-associated effects of glucocorticoid on BACE1 upregulation and Aβ generation: involvement of lipid raft-mediated CREB activation. Journal of Neuroscience 37: 8459–8476

Davidkova G, Carroll RC (2007) Characterization of the role of microtubule-associated protein 1B in metabotropic glutamate receptor-mediated endocytosis of AMPA receptors in hippocampus. Journal of Neuroscience 27: 13273–13278

Dent EW (2017) Of microtubules and memory: implications for microtubule dynamics in dendrites and spines. Molecular biology of the cell 28: 1–8

Du J, McEwen B, Manji HK (2009a) Glucocorticoid receptors modulate mitochondrial function: A novel mechanism for neuroprotection. Communicative & integrative biology 2: 350–352

Du J, Wang Y, Hunter R, Wei Y, Blumenthal R, Falke C, Khairova R, Zhou R, Yuan P, Machado-Vieira R (2009b) Dynamic regulation of mitochondrial function by glucocorticoids. Proceedings of the National Academy of Sciences 106: 3543–3548

Fellner S, Bauer B, Miller DS, Schaffrik M, Fankhänel M, Spruß T, Bernhardt G, Graeff C, Färber L, Gschaidmeier H (2002) Transport of paclitaxel (Taxol) across the blood-brain barrier in vitro and in vivo. The Journal of clinical investigation 110: 1309–1318

Friedman JR, Webster BM, Mastronarde DN, Verhey KJ, Voeltz GK (2010) ER sliding dynamics and ER– mitochondrial contacts occur on acetylated microtubules. The Journal of cell biology 190: 363–375

Galluzzi L, Pietrocola F, Levine B, Kroemer G (2014) Metabolic control of autophagy. Cell 159: 1263–1276

Gomez-Suaga P, Paillusson S, Stoica R, Noble W, Hanger DP, Miller CC (2017) The ER-mitochondria tethering complex VAPB-PTPIP51 regulates autophagy. Current Biology 27: 371–385

Green KN, Billings LM, Roozendaal B, McGaugh JL, LaFerla FM (2006) Glucocorticoids increase amyloid-β and tau pathology in a mouse model of Alzheimer’s disease. Journal of Neuroscience 26: 9047–9056

Grenningloh G, Soehrman S, Bondallaz P, Ruchti E, Cadas H (2004) Role of the microtubule destabilizing proteins SCG10 and stathmin in neuronal growth. Developmental Neurobiology 58: 60–69

Gustafsson N, Culley S, Ashdown G, Owen DM, Pereira PM, Henriques R (2016) Fast live-cell conventional fluorophore nanoscopy with ImageJ through super-resolution radial fluctuations. Nature communications 7: 12471

He M, Ding Y, Chu C, Tang J, Xiao Q, Luo Z-G (2016) Autophagy induction stabilizes microtubules and promotes axon regeneration after spinal cord injury. Proceedings of the National Academy of Sciences 113: 11324–11329

Hedskog L, Pinho CM, Filadi R, Rönnbäck A, Hertwig L, Wiehager B, Larssen P, Gellhaar S, Sandebring A, Westerlund M (2013) Modulation of the endoplasmic reticulum–mitochondria interface in Alzheimer’s disease and related models. Proceedings of the National Academy of Sciences 110: 7916–7921

Hunter RG, Seligsohn Ma, Rubin TG, Griffiths BB, Ozdemir Y, Pfaff DW, Datson NA, McEwen BS (2016) Stress and corticosteroids regulate rat hippocampal mitochondrial DNA gene expression via the glucocorticoid receptor. Proceedings of the National Academy of Sciences 113: 9099–9104

Kaganovsky K, Wang CY (2016) How do microtubule dynamics relate to the hallmarks of learning and memory? Journal of Neuroscience 36: 5911–5913

Kershaw S, Morgan D, Poolman T, Brass A, Matthews L, Ray D (2015) Glucocorticoids stabilise the microtubule network to inhibit cell migration. Endocrine abstracts 38 OC3.2

Krugers HJ, Hoogenraad CC, Groc L (2010) Stress hormones and AMPA receptor trafficking in synaptic plasticity and memory. Nature Reviews Neuroscience 11: 675

Lu W-Y, Man H-Y, Ju W, Trimble WS, MacDonald JF, Wang YT (2001) Activation of synaptic NMDA receptors induces membrane insertion of new AMPA receptors and LTP in cultured hippocampal neurons. Neuron 29: 243–254

Malagelada C, Jin ZH, Jackson-Lewis V, Przedborski S, Greene LA (2010) Rapamycin protects against neuron death in in vitro and in vivo models of Parkinson’s disease. Journal of Neuroscience 30: 1166–1175

Mandal M, Wei J, Zhong P, Cheng J, Duffney LJ, Liu W, Yuen EY, Twelvetrees AE, Li S, Li X-J (2011) Impaired α-amino-3-hydroxy-5-methyl-4-isoxazolepropionic acid (AMPA) receptor trafficking and function by mutant huntingtin. Journal of Biological Chemistry 286: 33719–33728

McEwen BS, Bowles NP, Gray JD, Hill MN, Hunter RG, Karatsoreos IN, Nasca C (2015) Mechanisms of stress in the brain. Nature neuroscience 18: 1353

Molitoris JK, McColl KS, Swerdlow S, Matsuyama M, Lam M, Finkel TH, Matsuyama S, Distelhorst CW (2011) Glucocorticoid elevation of dexamethasone-induced gene 2 (Dig2/RTP801/REDD1) protein mediates autophagy in lymphocytes. Journal of Biological Chemistry 286: 30181–30189

Nunan J, Shearman MS, Checler F, Cappai R, Evin G, Beyreuther K, Masters CL, Small DH (2001) The C◻terminal fragment of the Alzheimer’s disease amyloid protein precursor is degraded by a proteasome◻dependent mechanism distinct from γ◻secretase. European Journal of Biochemistry 268: 5329–5336

Olmos-Alonso A, Schetters ST, Sri S, Askew K, Mancuso R, Vargas-Caballero M, Holscher C, Perry VH, Gomez-Nicola D (2016) Pharmacological targeting of CSF1R inhibits microglial proliferation and prevents the progression of Alzheimer’s-like pathology. Brain 139: 891–907

Paillusson S, Stoica R, Gomez-Suaga P, Lau DH, Mueller S, Miller T, Miller CC (2016) There’s something wrong with my MAM; the ER–mitochondria axis and neurodegenerative diseases. Trends in neurosciences 39: 146–157

Pera M, Larrea D, Guardia◻Laguarta C, Montesinos J, Velasco KR, Agrawal RR, Xu Y, Chan RB, Di Paolo G, Mehler MF (2017) Increased localization of APP◻C99 in mitochondria◻associated ER membranes causes mitochondrial dysfunction in Alzheimer disease. The EMBO journal 36: 3356–3371

Picard M, McEwen BS (2014) Mitochondria impact brain function and cognition. Proceedings of the National Academy of Sciences 111: 7–8

Pinho CM, Teixeira PF, Glaser E (2014) Mitochondrial import and degradation of amyloid-β peptide. Biochimica et Biophysica Acta (BBA)-Bioenergetics 1837: 1069–1074

Psarra A-MG, Sekeris CE (2011) Glucocorticoids induce mitochondrial gene transcription in HepG2 cells: role of the mitochondrial glucocorticoid receptor. Biochimica et Biophysica Acta (BBA)-Molecular Cell Research 1813: 1814–1821

Real PJ, Tosello V, Palomero T, Castillo M, Hernando E, De Stanchina E, Sulis ML, Barnes K, Sawai C, Homminga I (2009) γ-secretase inhibitors reverse glucocorticoid resistance in T cell acute lymphoblastic leukemia. Nature medicine 15: 50

Schon EA, Area-Gomez E (2013) Mitochondria-associated ER membranes in Alzheimer disease. Molecular and Cellular Neuroscience 55: 26–36

Schreiner B, Hedskog L, Wiehager B, Ankarcrona M (2015) Amyloid-β peptides are generated in mitochondria-associated endoplasmic reticulum membranes. Journal of Alzheimer’s Disease 43: 369–374

Shaid S, Brandts C, Serve H, Dikic I (2013) Ubiquitination and selective autophagy. Cell death and differentiation 20: 21

Shen G, Ren H, Shang Q, Qiu T, Yu X, Zhang Z, Huang J, Zhao W, Zhang Y, Liang D (2018) Autophagy as a target for glucocorticoid-induced osteoporosis therapy. Cellular and Molecular Life Sciences: 1–11

Shin JE, Geisler S, DiAntonio A (2014) Dynamic regulation of SCG10 in regenerating axons after injury. Experimental neurology 252: 1–11

Song W-H, Yi Y-J, Sutovsky M, Meyers S, Sutovsky P (2016) Autophagy and ubiquitin–proteasome system contribute to sperm mitophagy after mammalian fertilization. Proceedings of the National Academy of Sciences 113: E5261–E5270

Sotiropoulos I, Silva J, Kimura T, Rodrigues AJ, Costa P, Almeida OF, Sousa N, Takashima A (2015) Female hippocampus vulnerability to environmental stress, a precipitating factor in tau aggregation pathology. Journal of Alzheimer’s Disease 43: 763–774

Uchida S, Martel G, Pavlowsky A, Takizawa S, Hevi C, Watanabe Y, Kandel ER, Alarcon JM, Shumyatsky GP (2014) Learning-induced and stathmin-dependent changes in microtubule stability are critical for memory and disrupted in ageing. Nature communications 5: 4389

Uchida S, Shumyatsky GP (2015) Deceivingly dynamic: learning-dependent changes in stathmin and microtubules. Neurobiology of learning and memory 124: 52–61

Vaux DL (2012) Research methods: Know when your numbers are significant. Nature 492: 180

Vershinin M, Carter BC, Razafsky DS, King SJ, Gross SP (2007) Multiple-motor based transport and its regulation by Tau. Proceedings of the National Academy of Sciences 104: 87–92

Wang J, Wang R, Wang H, Yang X, Yang J, Xiong W, Wen Q, Ma L (2017) Glucocorticoids suppress antimicrobial autophagy and nitric oxide production and facilitate mycobacterial survival in macrophages. Scientific reports 7: 982

Yao J, Irwin RW, Zhao L, Nilsen J, Hamilton RT, Brinton RD (2009) Mitochondrial bioenergetic deficit precedes Alzheimer’s pathology in female mouse model of Alzheimer’s disease. Proceedings of the National Academy of Sciences 106: 14670–14675

